# Bacterial flagella disrupt host cell membranes and interact with cytoskeletal components

**DOI:** 10.1101/2020.02.12.945204

**Authors:** Eliza. B. Wolfson, Johanna Elvidge, Amin Tahoun, Trudi Gillespie, Judith Mantell, Sean P. McAteer, Yannick Rossez, Edith Paxton, Fiona Lane, Darren J. Shaw, Andrew C. Gill, Jo Stevens, Paul Verkade, Ariel Blocker, Arvind Mahajan, David L. Gally

**Affiliations:** Divisions of Immunity and Infection, The Roslin Institute and R(D)SVS, The University of Edinburgh, Easter Bush, Midlothian, EH25 9RG, UK; Faculty of Veterinary Medicine, Kafrelsheikh University, 33516 Kafr el-Sheikh, Egypt; IMPACT Facility, Centre for Integrative Physiology, University of Edinburgh, Edinburgh, EH8 9XD, UK; Departments of Biochemistry, Biomedical Sciences Building, The University of Bristol, Bristol, BS8 1TD; Génie Enzymatique et Cellulaire, UMR 7025 CNRS, Centre de recherche Royallieu, Sorbonne Universités, Université de Technologie de Compiègne, Compiègne Cedex, France; Neurobiology, The Roslin Institute and R(D)SVS, The University of Edinburgh, Easter Bush, Midlothian, EH25 9RG, UK; Clinical Sciences, The Roslin Institute and R(D)SVS, The University of Edinburgh, Easter Bush, Midlothian, EH25 9RG, UK; Cellular and Molecular Medicine, Biomedical Sciences Building, The University of Bristol, Bristol, BS8 1TD

**Keywords:** Flagella, adherence, Escherichia coli, Salmonella, membrane, cytoskeleton, actin

## Abstract

Bacterial flagella have many established roles beyond swimming motility. Despite clear evidence of flagella-dependent adherence, the specificity of ligands and mechanisms of binding are still debated. In this study, the molecular basis of *E. coli* O157:H7 and *S*. Typhimurium flagella binding to epithelial cell cultures was investigated. Flagella interactions with host cell surfaces were intimate and crossed cellular boundaries as demarcated by actin and membrane labelling. SEM revealed flagella disappearing into cellular surfaces and TEM of *S*. Typhiumurium indicated host membrane deformation and disruption in proximity to flagella. Motor mutants of *E. coli* O157:H7 and *S*. Typhimurium caused reduced haemolysis compared to wild-type, indicating that membrane disruption was in part due to flagella rotation. Flagella from *E. coli* O157 (H7), EPEC O127 (H6), and *S*. Typhimurium (P1 & P2 flagella) were shown to bind to purified intracellular components of the actin cytoskeleton and directly increase in vitro actin polymerisation rates.

## Introduction

Bacterial flagella are protein organelles predominately associated with movement toward preferred environmental niches (1). They are comprised of a long, hollow, capped filament of polymeric flagellin, connected to a flagella basal body through a hook complex (2). The basal body houses a motor complex that rotates the basal body rod, hook and semi-rigid helical flagella filament, or flagellum, to move the bacterium (3, 4). The basal body also contains an adapted type 3 secretion export system that regulates secretion of the different organelle components during flagellum assembly (5, 6). Flagellin monomers make up the majority of the flagellum and have a hairpin-like structure with highly conserved associating termini and central variable domains (7). *Salmonella enterica* serovar Typhimurium can alternately express two flagella types, presumably to help avoid the adaptive immune response; phase-1 (P1) flagella are composed of FliC flagellin, and phase-2 (P2) flagella are composed of FljB flagellin (8). In the stacked flagella filament, the flagellin region exposed on the surface shows little conservation between strains and is used for immunological serotyping, resulting in the ‘H-type’. The flagellin termini are stacked inside the filament and drive flagellin polymerisation (9). These more conserved and buried regions are important microbe-associated molecular patterns (MAMPs), which activate Toll-like receptor 5 (TLR5) and NOD-like and CARD domain containing receptor 4 (NLRC4) (10–13).

In peritrichous and monotrichous species, flagella have been shown to sense surfaces and cause a switch from motile to sessile lifestyles, making flagellar adhesion the preliminary step essential for bacterial colonisation (14). More recently, flagellar rotation has been shown to play a role in flagella adherence, surface stiffness sensing and inducing Neutrophil extracellular traps (15, 16). However, mechanistic information beyond near-surface swimming is lacking and direct adherence along the filament shaft to cell surface components is poorly defined.

Important zoonotic enteropathogens, enterohaemorrhagic *E. coli* O157:H7 and *Salmonella enterica* serovar Typhimurium both express flagella and type 3 secretion systems (T3SSs) but have different intestinal colonisation strategies. T3SSs are structurally related to the flagella basal body and act as molecular needles to inject effector proteins into host cells (17). *E. coli* O157:H7 use their T3SSs to inject effector proteins into host intestinal epithelial cells, inducing actin polymerisation and the formation of attaching and effacing lesions (18). This cytoskeletal network modulation allows *E. coli* O157 to bind very tightly to the surface of host enterocytes while avoiding phagocytosis (19, 20). In contrast to *E. coli* O157:H7, *S*. Typhimurium translocates T3SS effectors into M-cells, inducing the rearrangement of the actin cytoskeleton into membrane ruffles, leading to bacterial uptake by M-cells or intestinal macrophages (21). *E. coli* O157:H7 is mainly an extracellular bacterium, but is reliant on both a T3SS and its H7 flagella for cell binding and colonisation of its reservoir host, cattle (22–24). Published studies indicate that H7 flagella bind to mucus and more recent work demonstrated binding of different *E. coli* flagella, including H7, to ionic phospho- and sulpho-lipids (25, 26). In contrast, *S.* Typhimurium is invasive in humans and animals, including cattle and pigs, with invasion being dependent on two different T3SSs (27). However, the significance of flagella expression by *S*. Typhimurium for initial attachment and the identity of potential protein ligands remain inconclusive (28–30).

This study takes a closer look at the molecular basis of bacterial flagella adherence to host cells in the context of pathogenic colonisation. We start by looking at actin associated with bacterial flagella when bacteria are in contact with host cells. We observe deformation and disruption of host plasma membranes in proximity to flagella and explore possible consequences of membrane breach. Actin and flagella interactions were confirmed by pull downs, far-Western blotting and *in vitro* actin polymerisation assays. These data shed some light on the interaction between *E. coli* O157:H7 and *S*. Typhimurium flagella and host cells and point towards the possibility of actin rearrangement and membrane ruffling induced by bacterial flagella.

## Results

### Bacterial flagella labelling can be coincident with host F-actin staining

In experimental systems that allow bacteria to come into contact with host cells without centrifugation, co-incidence of flagella and F-actin labelling was routinely detected, 60 minutes post-infection for *E. coli* O157:H7 (Fig. 1A-B, Fig. S1 & Movie S1) and 20 minutes post-infection for *S.* Typhimurium (Fig. 1C). For *Salmonella,* co-incidence of flagella with F-actin staining implied that flagella were not necessarily confined to the *Salmonella* containing vacuole (SCV) during invasion. It was also possible to observe flagella expressed by extracellular *Salmonella* co-incident with actin at early infection time points. However, not all flagella that were imaged (bacterially associated or not) were found to be coincident with phalloidin-stained actin (Fig. 1D). Where flagella were coincident with F-actin staining, this followed a striated pattern, consistent with the helical wave of flagella filaments.

**Figure 1.**
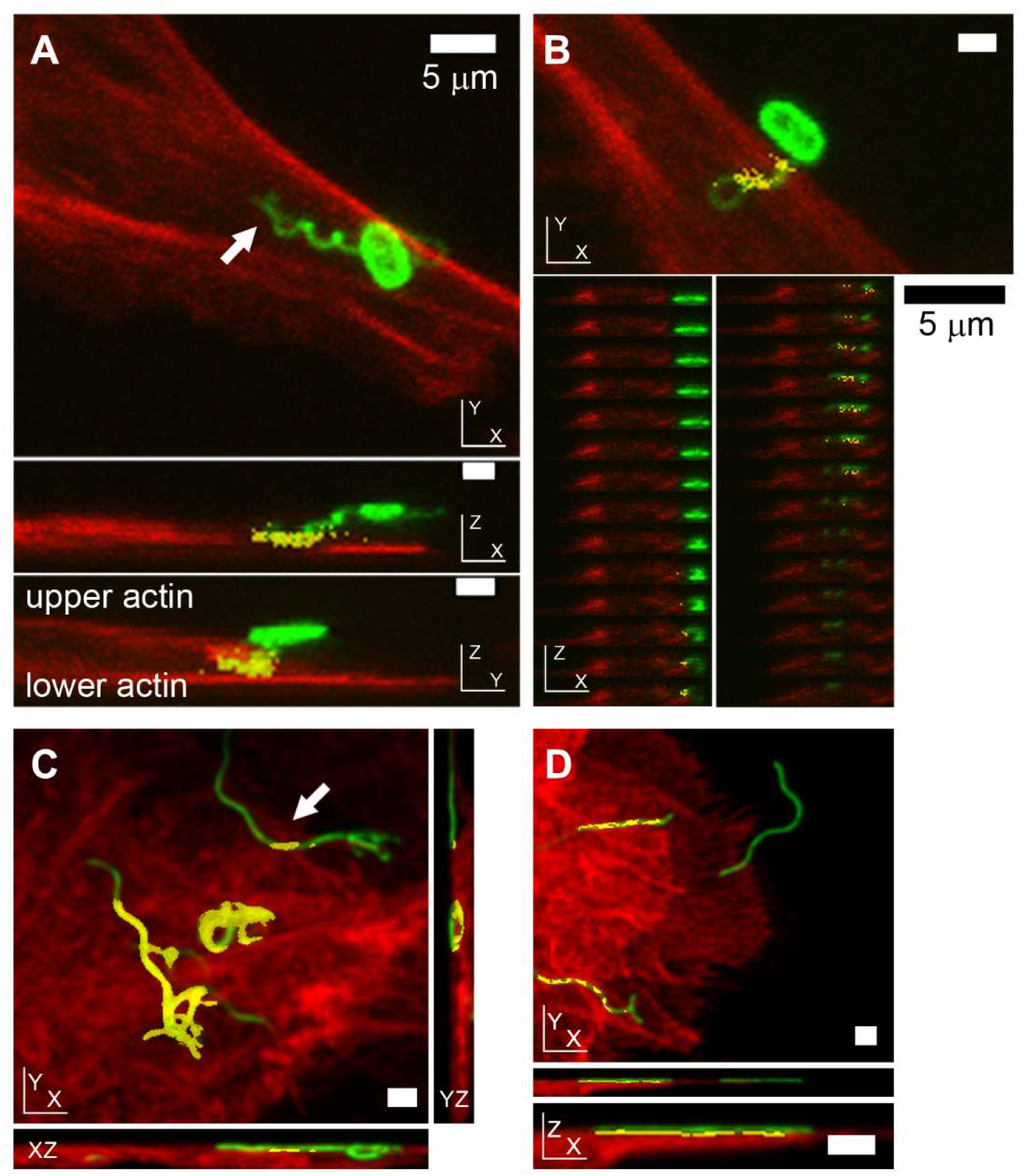
Position of *E. coli* O157:H7 and *S*. Typhimurium flagella relative to host cell F-actin during early colonisation by confocal microscopy. Phalloidin-labelled actin is red, antibody-labelled LPS and flagella are green, coincidence of the two is yellow. **(A-B)** Confocal micrographs of *E. coli* O157:H7 (TUV93-0) interacting with primary bovine epithelial cell cultures, 1 h post-infection. **(A)** Interaction of *E. coli* O157:H7 with host actin with its H7 flagellum in XY, XZ and YZ projections (3D rotating image in Movie S1, individual Z-slices of XZ and YZ projections are presented in Fig. S1). **(B)** Co-incidence of phalloidin-stained actin with an *E. coli* O157-associated coiled H7 flagellum, but not with the bacterium itself, by XY projection, with individual XZ slices shown beneath. **(C-D)** Confocal micrographs of O4:P2 stained *S.* Typhimurium (SL1344) interacting with IPEC-J2 cells, 20 min post infection. **(C)** XY and orthogonal projections of a 3D micrograph where a flagellum emanating from a non-invasive bacterium coincident with actin staining is indicated by an arrow. **(D)** XY projection of a 3D confocal micrograph of P2 flagella (green) adhering to the cell surface. Not all P2 flagella are coincident with actin, but where there is co-incidence, it is periodic. All samples were fixed with 4% (w/v) paraformaldehyde for >20 min prior to permeabilisation. Scale bars = 1 μm unless indicated.

### Bacterial flagella labelling is intermittently masked by membranes

Coincidence of bacterial flagella with F-actin staining was detected after fixation with 4% (w/v) PFA then permeabilisation with 0.1% (w/v) Triton X-100 (Tx100). To observe whether Tx100 treatment altered flagella labelling, confocal microscopy was carried out on GFP-expressing *E. coli* O157:H7 and *S.* Typhimurium SL1344 colonised epithelial cells. Cells were fixed and then flagella were labelled with specific antibodies before (red) and after (green) Tx100 treatment. F-actin was then labelled with AlexaFluor647 phalloidin (blue, Methods). Cell-associated flagellated *S.* Typhimurium and *E. coli* were imaged in 3D-stacks and representative projections are presented in figure 2. Pre-permeabilisation staining of flagella was often interrupted or less intense in short portions of the flagella, where post-permeabilisation staining of the same flagella was relatively even (Fig. 2). This indicates that bacterial flagella were intermittently masked by a detergent sensitive component during initial stages of infection.

**Figure 2.**
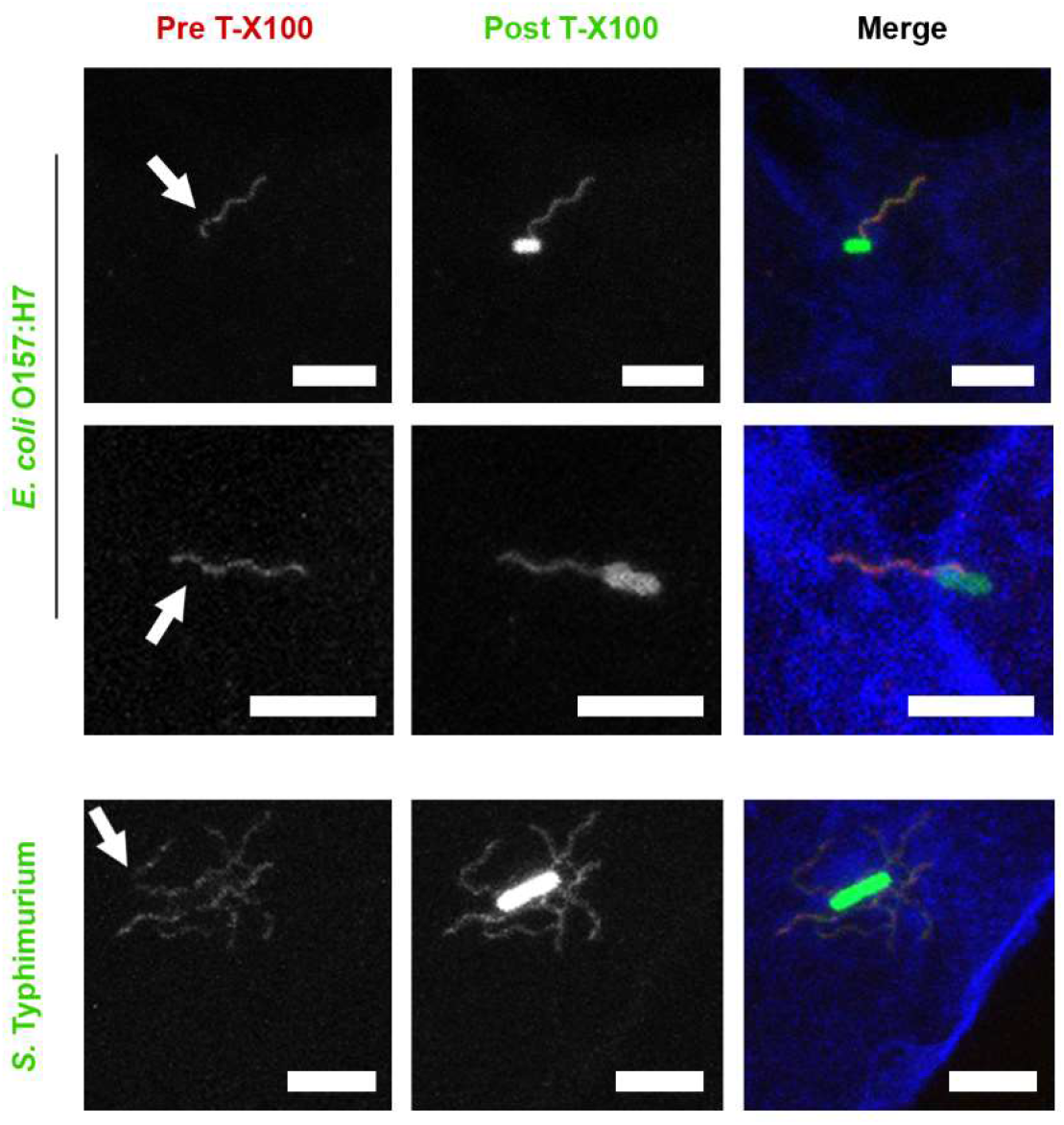
Confocal microscopy of *E. coli* O157:H7 and *S.* Typhimurium flagella staining before and after Triton X-100 treatment. *E. coli* O157:H7 TUV93-0∷pAJR145 (constitutively expressing GFP) interacting with EBL epithelial cells for 1 h, and *S.* Typhimurium SL1344∷pAJR145 interacting with IPEC-J2 epithelial cells for 20 min before fixation with 4% (w/v) paraformaldehyde for >20 min. Prior to permeabilisation, flagella were labelled with antibodies (red). After permeabilisation with 0.1% (v/v) Triton X-100 for 30 s, flagella were labelled with antibodies again (green). Cells were labelled with phalloidin-AlexaFluor647 (blue). Scale bars = 5 μm.

### Bacterial flagella are detectable on both sides of host cell membranes

To investigate how close flagella associations with host cell surfaces might be, flagellated adherent *E. coli* O157:H7 and *S.* Typhimurium were imaged in the context of plasma membrane labelling. For *E. coli* O157:H7 infections, HEK293 cells were transiently transfected with a trans-membrane voltage-gated ion channel fusion protein. This ion channel contains a green fluorescent protein (GFP) in the C-terminal cytoplasmic domain. The fluorescent signal generated reports the position of the internal cytoplasmic face of the plasma membrane only (31). H7 flagella were imaged passing in and out of the GFP-labelled cytoplasmic face of the plasma membrane (Fig. 3A, 3C). Volumetric 3D reconstruction of the labelling in these confocal microscopy images allowed the visualization of transverse sections, showing H7 flagella labelling within the cell cytoplasmic boundaries (Fig. 3B, 3D).

**Figure 3.**
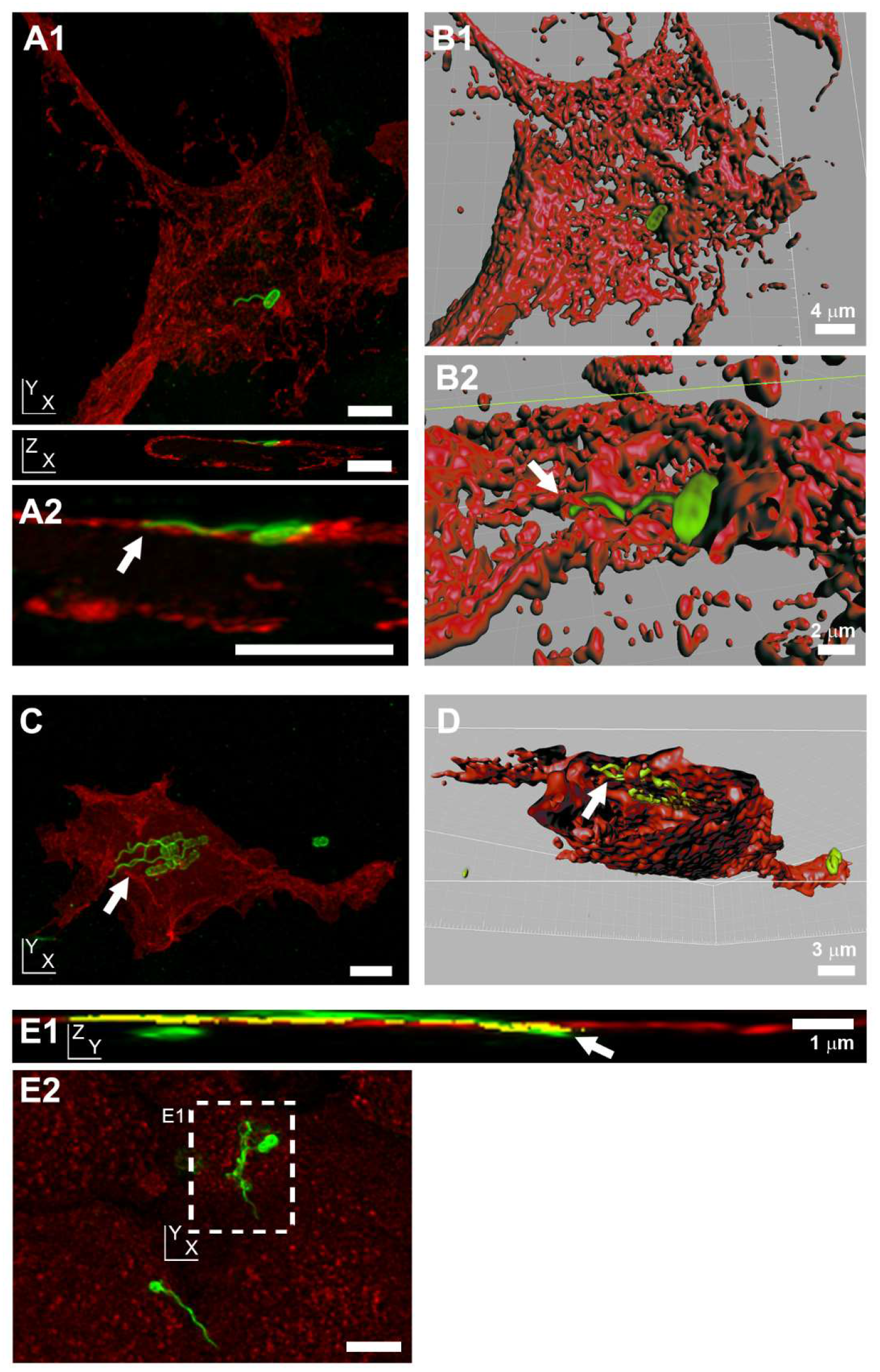
Position of *E. coli* O157:H7 and *S.* Typhimurium flagella relative to host cell membranes by confocal microscopy. **(A and C)** Projections of 3D confocal micrographs of *E. coli* O157:H7 (TUV93-0: LPS and flagella are green) interacting with HEK293 cells transiently transfected with a transmembrane protein-fluorescent protein fusion construct (red, Materials and Methods) 1.5 h post infection. This transfection causes labelling of the cytoplasmic face of the plasma membrane cytosolic leaflet (31). **(B and D)** 3D volume rendered model of images in (A) and (C) respectively, using Velocity Visualisation software. Arrows indicate intracellular flagella. **(A1)** XY and XZ projections of a single flagellum emanating from a bacterium on a transfected HEK293 cell, 3D rendered in **(B1)** and enlarged in **(A2)**. A similar view has been 3D rendered in **(B2)**, where the arrow indicates intracellular flagella. **(C)** XY projection of the binding of a cluster of bacteria expressing H7 flagella to a transfected HEK293 cell. **(D)** Transverse cut-through of the HEK293 cell, with flagella visible beneath the layer defined by the cytoplasmic staining of the labelled ion channel protein (arrow). **(E)** Projections of a 3D confocal micrograph of O4:P2 stained *S.* Typhimurium SL1344 (green) interacting with wheat-germ agglutinin labelled IPEC-J2 cells (red), 10 min post-infection. This labelling stains N-acetylglucosamine and sialic acid moieties on the external face of cell membranes (32). **(E1)** YZ projection of the XY inset labelled in (E2), with a long flagella bundle that passes out then back into the cell (arrow). Coincidence of flagella and membrane staining is shown in yellow. **(E2)** XY projection of whole image shown in (E1). All samples were fixed in 4% (w/v) para-formaldehyde for >20 min prior to permeabilisation. Scale bars = 5 μm unless indicated.

In contrast, wheat-germ agglutinin binds to the external sugar coated surface of epithelial cell membranes (32). Confocal microscopy of eukaryotic plasma membranes, labelled with fluorescent wheat-germ agglutinin after *S.* Typhimurium infection but before fixation, revealed flagella bundles of invasive *S.* Typhimurium passing through regions of plasma membrane staining (Fig. 3E). Flagella were observed passing both from inside-to-outside and outside-to-inside host cell areas. Both *E. coli* O157:H7 and *S.* Typhimurium flagella were observed passing through cellular boundaries by confocal microscopy.

### Defining bacterial flagella interactions with host cell membranes by electron microscopy

The z-plane resolution limit of confocal laser scanning microscopy is ~500 nm with the fluorophores used, but host cell plasma membranes are only ~5 nm thick. O157:H7 immuno-gold scanning electron microscopy (SEM) was used to take a closer look at H7 flagella interactions with host cell surfaces without Tx100 treatment. As with confocal microscopy of pre-permeabilisation labelling, *E. coli* O157:H7 flagella immune-gold staining was interrupted as the filament disappeared into the primary intestinal epithelial cell surface, then resumed as it curled upwards away from the surface (Fig. 4A-B). SEM of *S.* Typhimurium at early time points of infection (10-30 min, again without Tx100 treatment), also revealed flagella-like filaments interacting intimately with cell surfaces of primary intestinal epithelial cells by SEM (Fig. 4C). These filaments could be seen disappearing and reappearing from the cell surface, consistent with actin coincidence observed by confocal microscopy. *S.* Typhimurium flagella-like filaments were also observed associating with membrane ruffles and protrusions during the invasion process (Fig. 4D-E).

**Figure 4.**
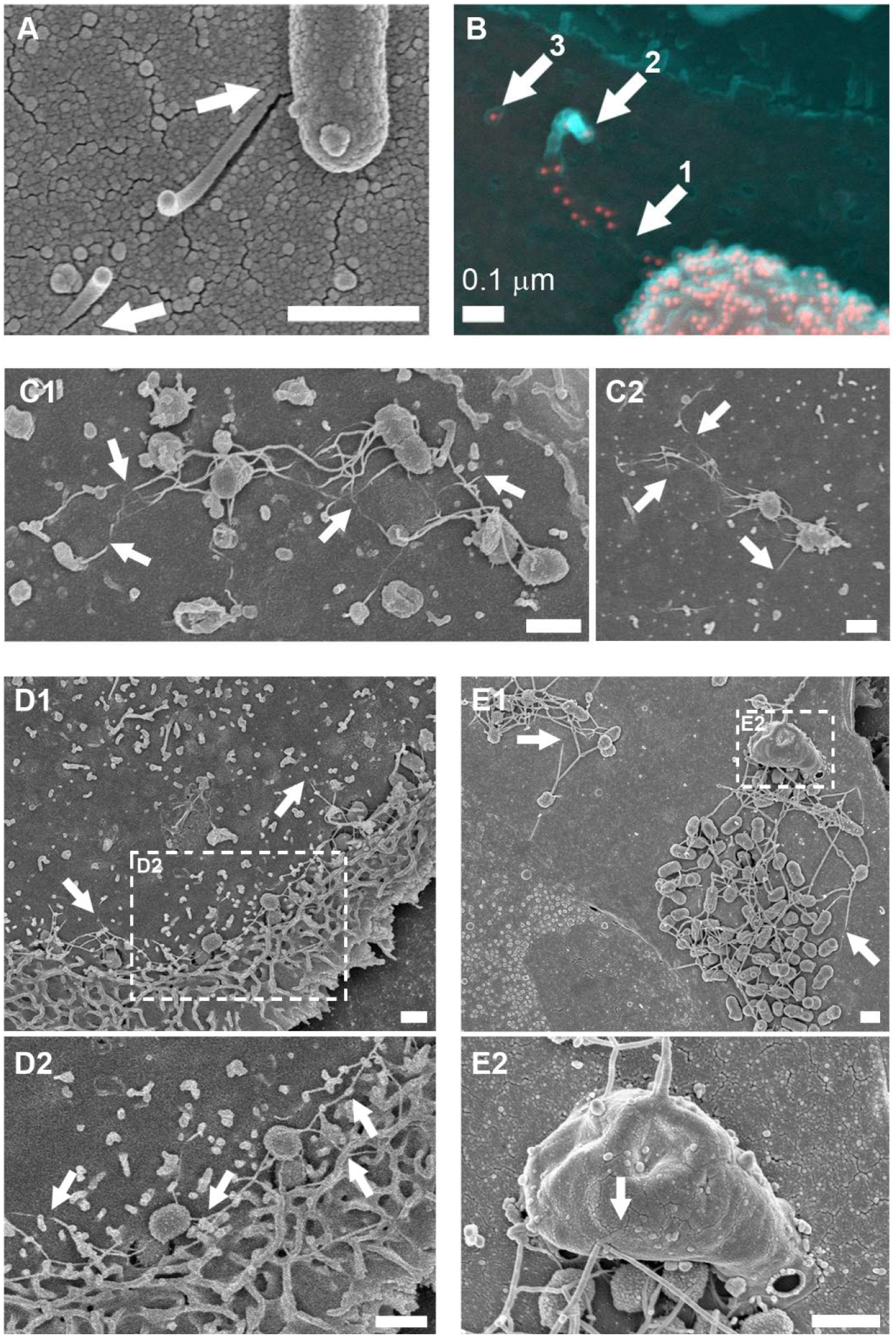
Scanning electron microscopy of *E. coli* O157:H7 and *S.* Typhimurium flagella-like structures disappearing and reappearing at host cell surfaces. **(A-B)** *E. coli* O157:H7 (ZAP734) interactions with bovine primary epithelial cell cultures, 3 h post-infection. **(A)** Secondary electron (SE) image of a snapped flagella-like filament, which appears to penetrate the cell surface (arrows). **(B)** shows a false coloured SE image (cyan) superimposed with false coloured back-scattered image (red). H7 flagella and O157 LPS are both immuno-gold labelled. Filament staining occurs adjacent to the bacterium and is then absent (arrow 1). The filament is then broken and curls back on itself (arrow 2), with the remnant embedded filament (arrow 3). **(C1-C2)** SE images of flagella-like filaments disappearing into and coming out of the surface of IPEC-J2 epithelial cells (arrows), within the first thirty minutes of infection with *S.* Typhimurium (Maskan). **(D)** SE image of *S.* Typhimurium (Maskan) micro-colonies on IPEC-J2 cells, 40 min post-infection. **(D1)** shows actin ruffling proximal to invading *Salmonella*. **(D2)** is enlarged from the inset indicated in D1. Wavy flagella-like filaments are interacting with ruffled and un-ruffled cell surfaces (arrows). **(E)** SE image of *S.* Typhimurium SL1344 (WT) micro-colonies on bovine primary epithelial cells. **(E1)** shows long filaments disappearing into the cell surface (arrows). **(E2)** is a higher resolution image of the area indicated in B1, which shows long filaments interacting with a large macropinocytic protrusion (arrow). Samples were fixed in 3% (w/v) glutaraldehyde and were not permeabilised prior to sample processing for scanning electron microscopy. Imaging was undertaken on a Hitachi 4700 Field Emission Scanning Electron microscope. Scale bars = 1 μm unless indicated.

These observations were made with serologically and structurally distinct flagella (Fig. S2), hinting that a biophysical mechanism may be involved. To look for evidence of biophysical interactions of flagella with host cell membranes, *S.* Typhimurium flagella closely associated with host cells were located with correlative light and electron microscopy and examined by thick-section transmission electron tomography (Fig. 5). To preserve the details and visibility of the host cell membranes, conventional post section staining of flagella was not undertaken, but the visibility of the structures in the tomogram slices was enhanced by applying an anisotropic diffusion filter and a blue-orange ICB colour lookup table in Amira software.

**Figure 5.**
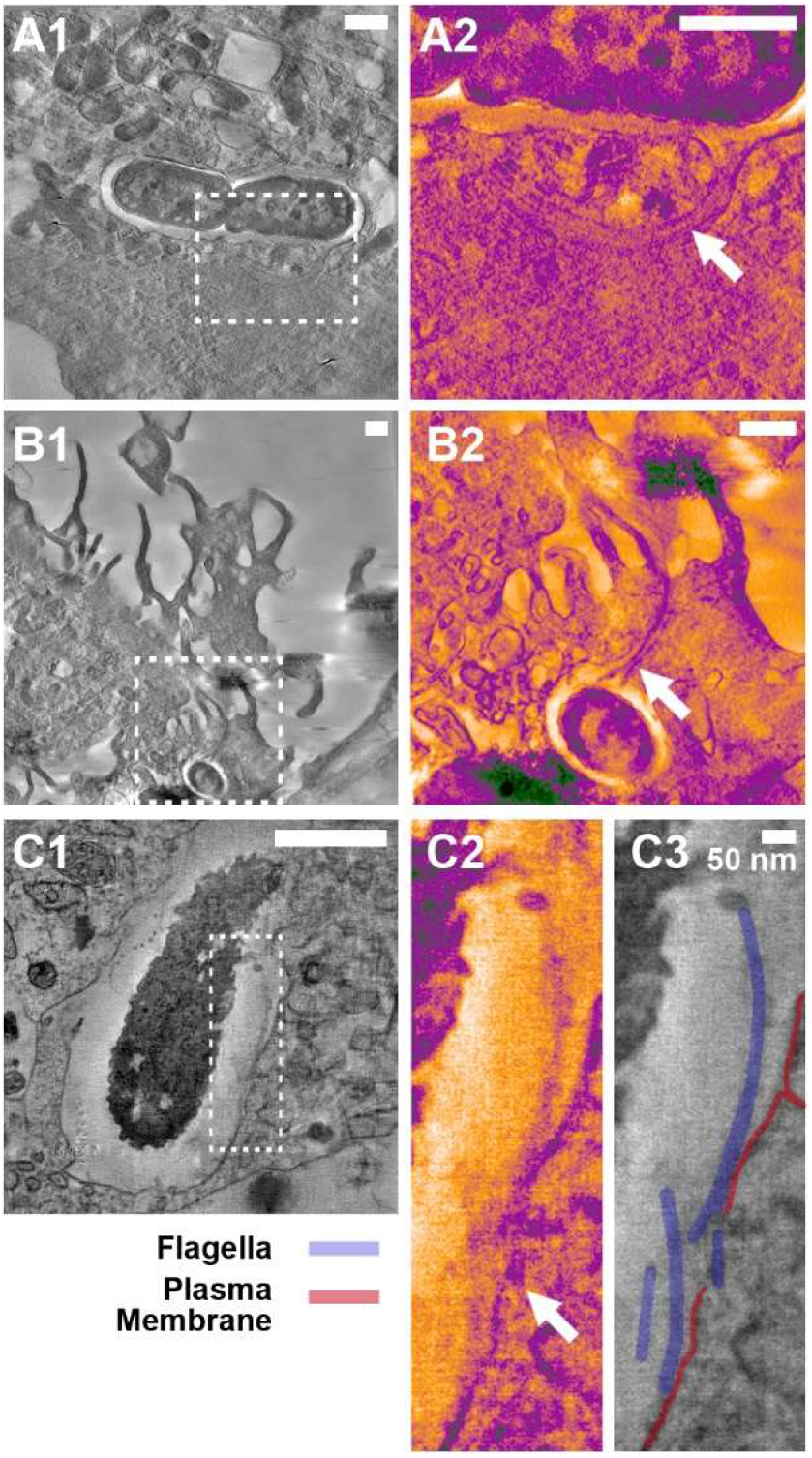
Electron tomography of host cell membranes in proximity to *S.* Typhimurium flagella-like structures. *S.* Typhimurium SL1344∷pAJR145 interacting with IPEC-J2 cells, 20 min post infection. Fixation with 4% (w/v) paraformaldehyde was undertaken for >20 min prior to flagella labelling and samples were not treated with any detergents. Confocal microscopy was used to locate bacteria with flagella in contact with host cells and correlative electron tomography was undertaken (Methods). The contrast of insets was enhanced using the blue_orange_ICB look up table in Fiji software. **(A1)** Flagellated *S.* Typhimurium within an epithelial cell. **(A2)** Inset in A1; the flagella cross the gross vacuole boundary but are present inside an intact membrane channel (arrow). **(B1)** Flagellated *S.* Typhimurium inside a membrane ruffle. **(B2)** Inset in B1; the flagella bundle passes through the gross boundaries of the ruffle but is adjacent to a distinct membrane boundary (arrow). **(C1)** Flagellated *S.* Typhimurium at the point of induced uptake into an epithelial cell. **(C2)** Inset in C1; flagella in proximity to the disrupted epithelial plasma membrane (arrow). **(C3)** Structures in C2 are labelled for clarity. Scale bars = 500 nm unless indicated.

Host cell membranes were observed encapsulating bundles of flagella filaments during the *S.* Typhimurium invasion process (Fig. 5A-B, Movies S2-3). Consistent with previous microscopy, these flagella were located within gross cellular boundaries at points, but they were separated from the host cytosol by deformed but contiguous host membranes. Visually distinct from these observations, was the reduction or interruption of clear membrane staining in proximity to bacterial flagella (Fig. 5C, Movie S4). In this example, the flagella filaments are largely accessible to extracellular antibody labelling, as shown by the punctae that occur regularly along their length. Figure 5C2 shows the labelled bundle of flagella filaments drawing close to a clearly defined plasma membrane, but as they come into presumable contact, the membrane boundary is both deformed and less distinct, and in places apparently absent, suggesting membrane disruption.

### Bacteria with paralysed flagella cause less membrane disruption

To test whether flagella rotation is capable of causing physical disruption of host cell membranes, sheep red blood cells (RBCs) were used as a model dye-filled eukaryotic plasma membrane, as they contain haem pigment. Short-term incubation of RBCs was undertaken with bacteria and their isogenic flagella and flagella motor (*mot)* mutants; *mot* mutants result in the production of full-length flagella that do not rotate. Bacteria were cultured under conditions known to induce expression of flagella and briefly centrifuged into contact with RBCs to eliminate the contribution of motility and chemotaxis to this process (Methods, Fig. 6).

**Figure 6.**
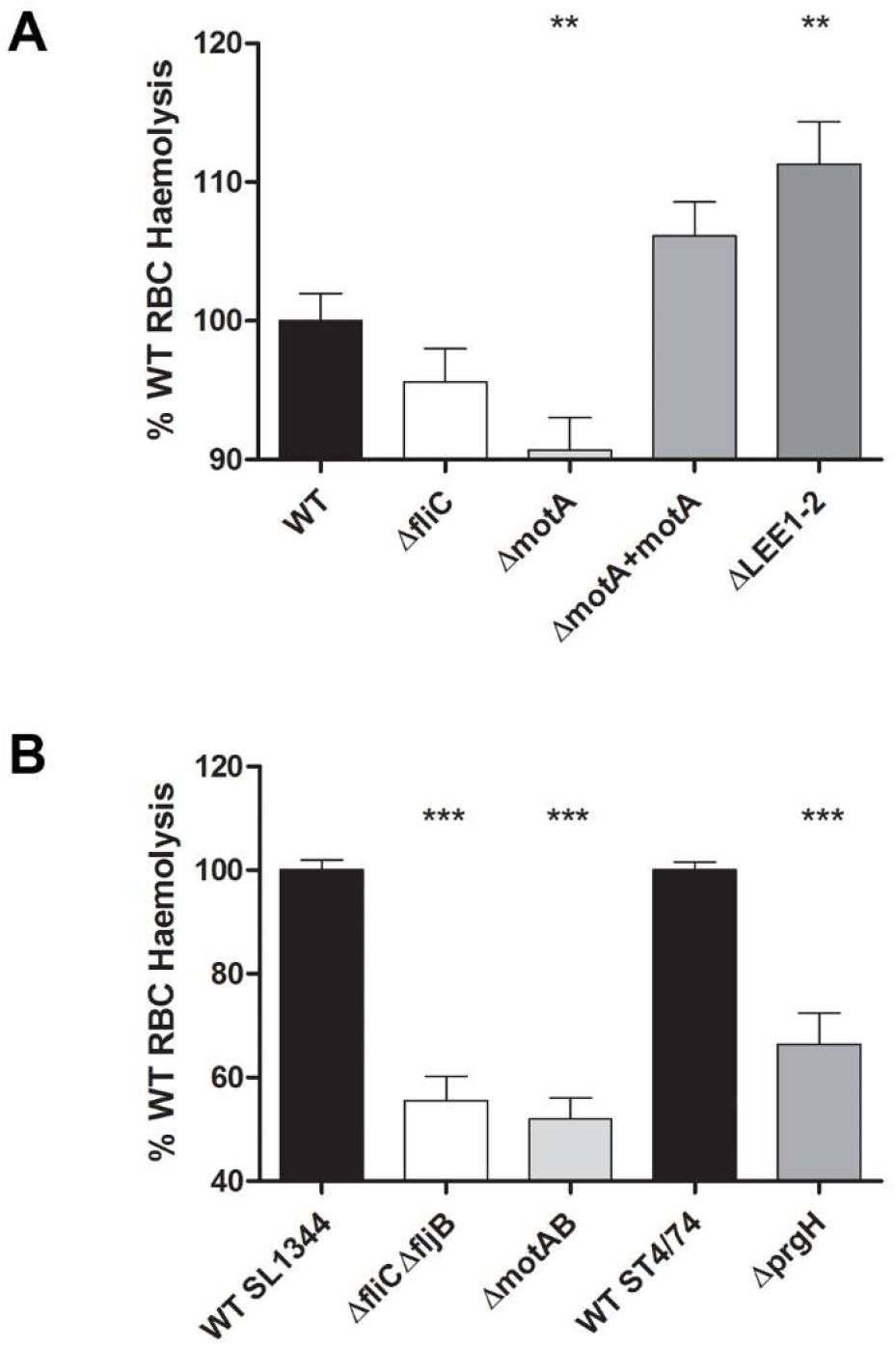
Contribution of flagella rotation to membrane disruption by *S*. Typhimurium and *E. coli* O157:H7. Sheep red blood cells (RBCs) were incubated with stationary phase cultures of the *E. coli* O157:H7 **(A)** and *S.* Typhimurium **(B)** strains indicated for 2 h at 37°C, after centrifugation together at 2000 × *g* for 5 min to ensure contact. After incubation, samples were centrifuged at 2000 × *g* for 10 min and the A_405_ of supernatants were measured. Data from 3-5 biological replicates are presented as % wild-type (WT) haemolysis and statistical analysis was performed on raw data using 2-tailed homoscedastic student’s t-tests; p≤0.01 (**), p≤0.001 (***). The results were considered as significant for a p value ≤0.05.

All flagellate wild-type strains caused membrane disruption upon interaction with RBCs, as determined by haem release. As haem release could be due to the T3SSs of either pathogen, T3SS mutants (ΔprgH for S. Typhimurium, ΔLEE1-2 for E. coli O157:H7, Table 1) were tested under these flagella-selective growth conditions (Fig. 6). Mutation of T3SSs resulted in altered mean levels of haem release compared to wild-type for both *S.* Typhimurium (~30% reduction) and *E. coli* O157:H7 (~10% increase). Both effects were statistically significant (p<0.0001 and 0.0039 respectively), suggesting that the T3SSs were directly involved in haem release.

**Table 1.** Strains used or constructed in this study.

| Strain | Relevant features | Source |
| --- | --- | --- |
| TUV93-0 | *E.coli* O157:H7; EDL933 (ATCC 700927) *stx*<sup>-</sup> | (62) |
| TUVΔ*fliC* | TUV93-0 *fliC*::*sacB*::*kan*<sup>r</sup>, aflagellate | (26) |
| TUV*fliC*<sup>-</sup> | TUV93-0 *fliC*<sup>-</sup>, aflagellate | (26) |
| TUV*fliC*<sub>H7F</sub> | TUVΔ*fliC* *cis* complement *fliC*<sub>H7</sub> from TUV93-0 | This study |
| TUV*fliC*<sub>H6F</sub> | TUVΔ*fliC* *cis* complement *fliC*<sub>H6</sub> from E2348/69 | This study |
| ZAP1574 | TUV93-0 Δ*motA* | This study |
| ZAP1575 | TUV93-0 *motA*<sup>-</sup> | This study |
| ZAP1576 | TUV93-0 *motA*-complemented | This study |
| ZAP734 | *E. coli* O157:H7 NCTC 12900, *Stx*<sup>-</sup> | AHVLA |
| E2348/69 | *E.coli* O127:H6; sequenced | (63) |
| MG1655 | *E. coli* K12; sequenced | (64) |
| TUV Δ*LEE1-2* | 8.9 Kb deletion in *LEE1-3*, *ETTA1* | (65) |
| SL1344 | *S. Typhimurium* | (66) |
| SL1344Δ*fljB*<sub>P2</sub> | SL1344 Δ*fljB* by λred, expresses only P1 | (66) |
| SL1344Δ*fliC*<sub>P1</sub> | SL1344 Δ*fliC* by λred, expresses only P2 | (66) |
| SL1344 Δ*fliC*<sub>P1</sub><br>Δ*fljB*<sub>P2</sub> | SL1344 Δ*fliC* and Δ*fljB* by λred, aflagellate | (66) |
| ZAP1566 | *S. Typhimurium* LT2 *fliM*5978-GFP<br>*motAB*::Tc<sup>R</sup>, non-motile | P. Aldridge Lab,<br>Newcastle |
| SL1344Δ*motAB* | SL1344 P22 transduced with ZAP1566, non-motile | This study |
| Maskan | *S. Typhimurium* | Stock from<br>Lohmann Animal<br>Health |
| 4/74 | *S. Typhimurium* | (67) |
| 4/74 *prgH*<sup>-</sup> | *prgH* : : mini-Tn 5Km2 mutant of ST4/74 NaI<sup>R</sup> | (68) |

Mutation of flagella and flagella rotation caused reduced mean levels of haem release for both *S*. Typhimurium and *E. coli* O157:H7. For *S*. Typhimurium, the levels of haem release were approximately half of wild-type in both flagellin (Δ*fliC*Δ*fljB*) and motility (Δ*motAB*) mutants, with statistically significant values (p<0.0001 for both, Fig. 6). For wild-type *E. coli* O157:H7, mean levels of haem release were much less affected by flagellin (*fliC^−^*) and motor (*motA^−^*) mutation. Reductions of ~5% for *fliC^−^* were not statistically significant (p=0.17). In contrast, a reduction of ~10% for *motA^−^* was statistically significant (p=0.0046). The subtle effect for *E. coli* O157:H7 may reflect the lower overall flagella expression, lower rotation speeds and motility of *E. coli* wild-type compared to *Salmonella* (Fig. S3, (33, 34)). These results suggest that the flagella of *S.* Typhimurium and *E. coli* O157:H7 are physically able to disrupt eukaryotic plasma membranes, but with different efficacies, and flagellar rotation is likely to play a key role.

### Bacterial flagella can interact with components of the actin cytoskeleton *in vitro*

If bacterial flagella can disrupt host cell membranes, perhaps they are also able to interact with cytosolic components just beneath host cell membranes. Initial pull downs and far-Western blotting with H7 flagella identified β-actin (ACTB1), cofilin-1 (CFL1) and galectin-4 (GAL4) as potential interactants (Fig. S4). A ‘far’ ELISA was designed to assess relative flagella binding to human βγ-actin, recombinant human CFL1 and recombinant human GAL4. These purified proteins were coated to 96-well plates, probed with 1000-100 ng flagellins, then binding was detected using appropriate antibodies. Binding data were normalized to negative (no flagellin added) and positive (wells coated with 1 μg flagellin) detection controls to give a measure of relative binding (Methods). Binding of polymeric and monomeric flagellin from *E. coli* O127:H6, *E. coli* O157:H7, commensal *E. coli* K12:O175:H48 and *S.* Typhimurium O4:Hi/H2 expressing either P1 or P2 flagella was assessed (Fig. 7, S2).

**Figure 7.**
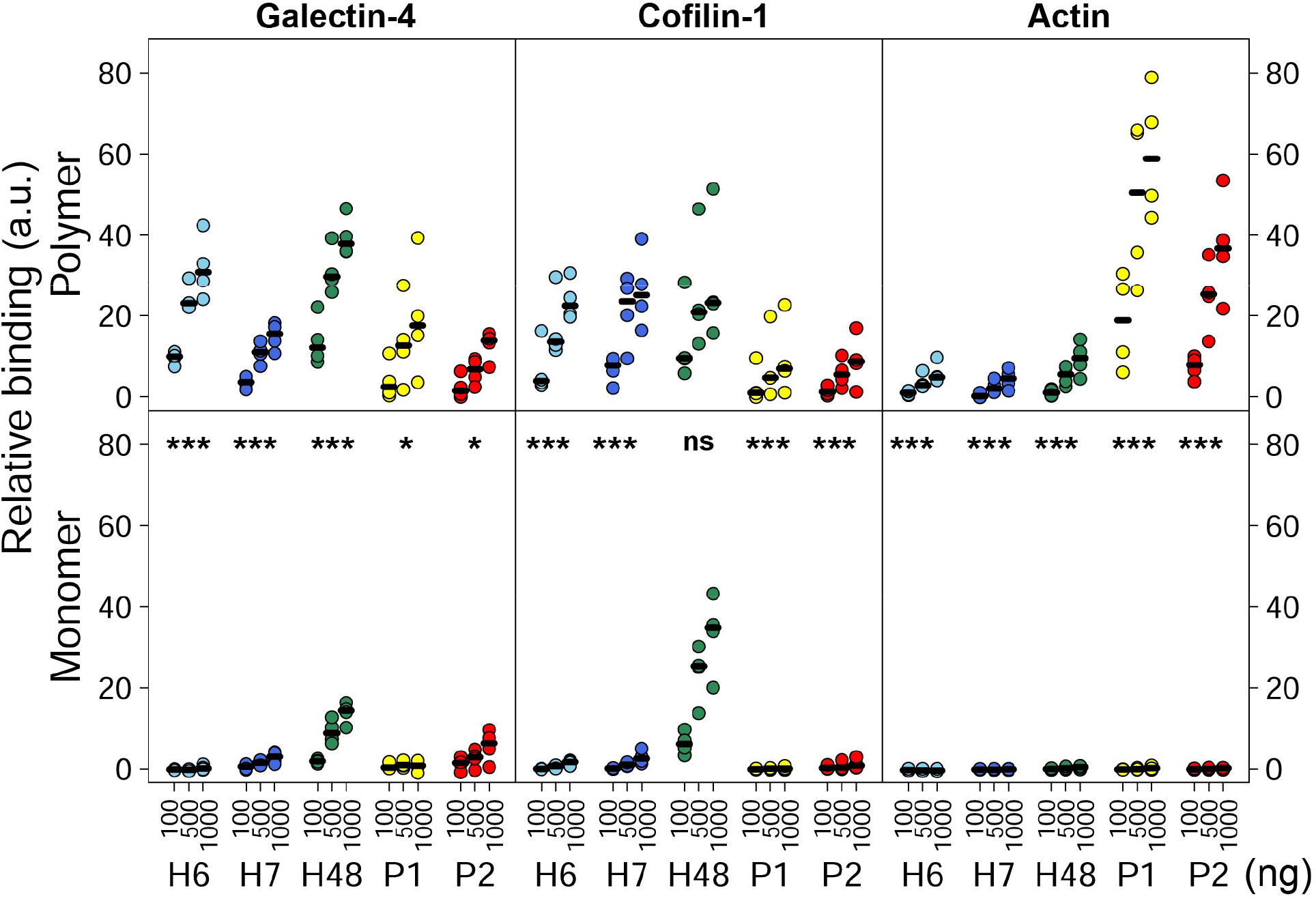
Relative binding of flagella from *E. coli* and *S.* Typhimurium to cytoskeletal proteins by far-ELISA. Flagella binding interactions with identified cellular components assessed by far-ELISA. Binding of 100, 500 and 1000 ng/well of polymeric and monomeric H6 (light blue), H7 (dark blue) and H48 flagella (green) from *E. coli*, and P1 (yellow) or P2 (red) flagella from *S.* Typhimurium to 1000 ng/well galectin-4, cofilin-1 and βγ-actin. Median values of four independent experiments are indicated with black horizontal lines. Relative binding levels are shown as data are normalised to positive and negative detection controls for each antibody and protein pair (Methods). Statistical analyses presented are pairwise comparisons of polymer and monomer using linear mixed effects models; ns, not significant, p≤0.01 (*), p≤0.0001(***).

For all interactions tested, except for H48-CFL1 (p=0.55), monomeric flagellins bound at significantly lower levels than polymeric flagellins (Fig. 7, p=0.0023-0.0001 or less, depending on the flagellin tested), serving as a suitable baseline for non-binding in this assay. This was particularly the case for the actin interactions, where binding by flagellin monomers was virtually undetectable in this assay for each flagellin type tested.

There were different patterns of binding between *E. coli* and *S.* Typhimurium polymeric flagellins; *E. coli* flagellins showed higher relative binding to CFL1 compared to βγ-actin under these conditions, and conversely the *S.* Typhimurium flagellins tested showed a stronger association with βγ-actin, compared to CFL1. All polymeric flagellins tested bound to GAL4, though there was no particular distinction between *S.* Typhimurium and *E. coli* relative levels. Binding to GAL4 was not a consequence of flagella post-translational glycosylation; determination of molecular weights by mass spectrometry of H6 and H7 sheared preparations indicated that these flagellins were unmodified, whilst modified forms of *S.* Typhimurium flagellins were consistent with methylation (data not shown, (35)).

### Bacterial flagella can increase actin polymerisation rates *in vitro*

To further confirm the *in vitro* binding interactions of H6, H7, P1 and P2 polymeric flagellin with actin, their effect on actin polymerisation was assessed (Fig. 8A). 1 mM pyrene-conjugated rabbit αβ-actin in 10 mM Tris-HCl, 200 μM CaCl_2_, 200 μM ATP, 1 mM DTT, pH 7.5, was used as a baseline (Actin control). Actin polymerisation was initiated by addition of final concentrations of 5 mM KCl, 0.2 mM MgCl_2_ and 0.1 mM ATP (Actin). The actin severing activity of CFL1 at 2:1 ratio with actin generates more rapidly polymerising ‘plus’ ends of actin (36), so 500 nM CFL1 was added as a positive control for enhanced rates of actin polymerisation (cofilin-1). The bacterial flagella filaments tested all enhanced *in vitro* rabbit αβ-actin polymerisation to variable extents, implying a direct interaction with physiologically active actin (Fig. 8A). Higher median actin polymerisation V_max_ rates were determined for all flagella tested when compared to actin alone, with P2 flagella showing a statistically significant dose-dependent effect (p=0.003, Fig 8A-B).

**Figure 8.**
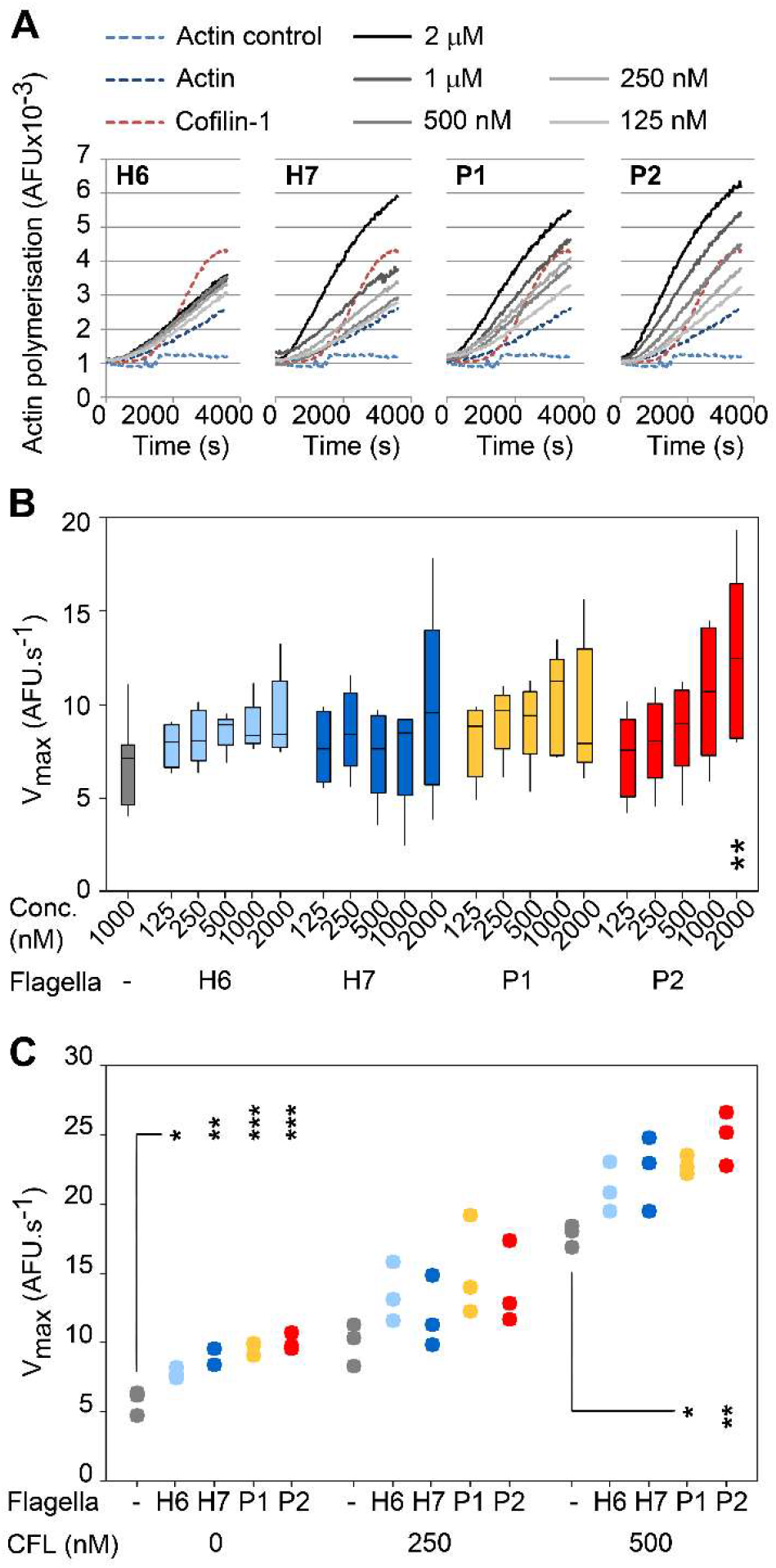
Impact of flagella type on actin polymerisation kinetics. **(A)** Effects of H6, H7, P1 and P2 flagella on αβ-actin polymerisation. Pyrene-conjugated globular (G)-actin polymerisation was triggered by addition of 5 mM KCl, 0.2 mM MgCl_2_ and 0.1 mM ATP in the presence of different concentrations of flagella, and arbitrary fluorescence (AFU) was measured every 30 s for 1 h at RT. 500 nM cofilin-1 (CFL) was used as a positive control for increased actin polymerisation rates in sub-optimal conditions (37). Data shown is a representative experiment. **(B)** Maximum velocity (V_max_) of actin polymerisation in the presence of titrating concentrations (125 nM − 2 μM) of H6 (light blue), H7 (dark blue), P1 (yellow) and P2 (red) flagella. Data was analysed from five independent experiments and post-hoc Tukey pairwise comparisons of differences between specific flagella concentrations and actin alone are presented; p<0.01 (**). **(C)** V_max_ of actin polymerisation in the presence of 250 nM H6, H7, P1 and P2 flagella, and sub-optimal (250 nM) and optimal (500 nM) concentrations of cofilin-1 (CFL). Data represents results from three independent experiments and post-hoc Tukey pairwise comparisons of the difference between CFL with and without different types of flagella are presented; p<0.05 (*), p≤0.01 (**), p≤0.001 (***).

To investigate whether H6 and H7 flagella interactions with CFL1 enhanced the effect they had on actin polymerisation, αβ-actin was co-incubated with the four flagella types and two different concentrations of CFL1 (Fig. 8C). Without CFL1, 250 nM of each flagella type was sufficient to cause a statistically significant increase in the V_max_ of actin polymerisation, with *S.* Typhiumurium P2 flagella being the most effective. This low concentration of flagella was chosen to allow detection of increases in actin polymerisation rates that may occur as a result of synergy between flagella and CFL1 interactions. Adding an equimolar concentration of CFL1 (250 nM) caused a trend towards further enhanced rates of actin polymerisation, but this was not statistically different to addition of CFL1 alone for any of the flagella types tested (p=0.124-0.856). At a 2:1 molar concentration of CFL1:flagella, at which the CFL1 concentration was optimal, further increases in the V_max_ of actin polymerisation were apparent for all flagella tested. These increases were only statistically significant with P1 and P2 flagella, compared to the effect of CFL1 alone (p=0.035 and 0.004 respectively). Additionally, there was no statistically significant interaction between CFL1 concentration and flagella type (p=0.6, Methods), indicating that the effects on V_max_ were additive and not due to any synergistic effects of flagella and CFL1 on actin polymerisation.

## Discussion

We have known for some time now that bacterial flagella are more than just motility organelles. The idea of flagella as a long adherence organelle for early host surface attachment by commensal and pathogenic bacteria alike is now widely accepted. This raises the question of why there are so few published examples of specific cell-surface protein receptors for flagella.

This paper presents the masking of *Salmonella* and *E. coli* flagella antigens by a detergent-sensitive component, which indicates very intimate interactions of these flagella with host cell lipids. Such a close interaction between flagella and membranes was borne out by confocal microscopy of membrane labelled samples. Additionally, flagella of *S.* Typhimurium and *E. coli* O157:H7 have been shown to bind to cholesterol and ionic sulpho- and phospho-lipids *in vitro* respectively (26, 38), so it is conceivable that these flagella could be binding directly to plasma membranes.

Several studies attribute flagella-mediated adherence to surfaces as largely biophysical processes. It has been demonstrated that *E. coli* H48 flagella are generally attracted to hydrophobic surfaces and are ten times more likely than the bacterial cell body to make contact with artificial surfaces. These collisions by flagella with hydrophobic abiotic surfaces are known to slow bacteria down, promoting adhesion (39, 40). This is also true in near surface swimming, a biophysical process dependent on rotating flagella, which has been shown to aid cooperative colonisation by probing and facilitating docking at membrane protrusions (41, 42). Additionally, the structural properties of different flagellin phase types have been shown to alter *S.* Typhimurium flagella rotation and near-surface swimming behaviour, with downstream effects on colonisation (43).

The observations of flagella crossing gross cellular boundaries presented here raise the possibility that flagella-mediated adherence may go beyond surface adhesion. This was routinely detected by confocal microscopy of actin and membrane labelled samples. Such overlaps could be explained by insufficient resolution of cellular boundaries, and electron tomography showed this to be at least partially the case. Flagella bundles attached to colonising bacteria were observed crossing gross cellular boundaries, but the majority remained enclosed within deformed membrane channels. This hints that biophysical membrane deformation by flagella may enhance host membrane ruffling, aiding flagella dependent adherence by creating favourable surface topologies for membrane binding. Where possible membrane disruption was observed, cells were fixed for microscopy, so there is the potential for observation artefacts resulting from sample dehydration. However, at the time of writing, the technology to obtain 3D sub-micrometre resolution of moving samples, labelled in a way that will not interfere with interacting surfaces, is not readily available. Nevertheless, the observations are supported by the additional phenotypic data that *E. coli* O157:H7 and *S.* Typhimurium cause 10% and 50% respectively less disruption of biological membranes without flagella rotation, compared to wild-type.

Where potential disruption of host cell membranes in proximity to bacterial flagella was observed by electron tomography, the mechanism by which they do this remains speculative. The haemolysis data supports a role for flagella rotation. Yet whether this membrane disruption is direct, due to biophysical force, or indirect, due to increased probabilities of favourably charged flagella colliding with membranes, is not known. Alternatively, on epithelial cells this process could be a consequence of pathogen-induced large-scale membrane rearrangements seen during *S.* Typhimurium and *E. coli* colonisation, trapping flagella at random, or directed by receptor-mediated endocytosis, triggered by flagella binding to relevant surface molecules. *P. aeruginosa* flagella bind to MUC1 and heparin-sulphated proteoglycans and *E. coli* flagella bind to ganglioside receptor GM1 and charged glycerolipids, resulting in immuno-modulation and invasion (26, 44–46). It is also possible that membrane penetration could be caused by a combination of these factors.

Interaction with actin cytoskeletal proteins was not restricted to flagella from pathogenic bacteria. As established, the predicted surface epitopes of the different polymerised flagellins studied are markedly distinct (Fig. S2), complicating analyses of key surface regions required for CFL1, galectin-4 and βγ-actin binding. Friedlander *et al.* (42) proposed that flagella binding to artificial surfaces involves low-affinity but high-avidity co-operative binding to surface substrates. Multiple low-affinity interactions of the different polymeric flagellins may also account for the direct binding to cytoskeletal proteins observed in the present study. This supports the concept that this co-operative binding of flagellin subunits within a filament is a generic property of flagella. In a pathogenic context this could lend itself to binding repetitive substrates like actin filaments. Flagella in a non-pathogenic context may be capable of binding to actin and actin binding proteins, but for this to be relevant, the flagella would have to gain access through the mucus layer to the epithelial surface. Conversely, motility and flagella-deficient strains of *E. coli* and *S.* Typhimurium can still be pathogenic (47, 48). However, where expressed, the importance of flagella in colonisation of enteropathogenic and enterohaemorrhagic *E. coli,* and *S.* Typhimurium has been established (22, 28, 43, 49), and this study highlights another potential reason for this.

Furthermore, a recent paper has identified a proteolytic site in the hypervariable region of flagella, present in at least 74 bacterial species, including several species of *Clostridium.* This metallopeptidase domain, named flagellinolysin was not found in either *E. coli* nor *S. enterica* (50). However, this raises the possibility that these flagella are capable of other enzymatic activities against actin or membrane lipids. Additionally, by using flagellinolysin negative species, this study could serve as a baseline in assessing the relative contribution of flagellinolysin to flagella-dependent colonisation processes in the future.

Flagella interactions with actin were undertaken with isolated components in physiological conditions, confirming direct binding. Many pathogens reorganise host actin architecture during host colonisation. The first bacteria described to use actin rearrangement were the intracellular bacteria *Listeria monocytogenes* and *Shigella flexneri* (51, 52). They both generate filamentous actin “comet tails” that propel the bacteria through the cytosol, allowing colonisation of neighbouring cells. Modulation of actin and actin binding proteins, such as Arp2/3 and CFL1 has also been described for certain pathogenic *E. coli* and *Salmonella* spp. secreted effector proteins (53, 54). However, the findings in this current study provide the first evidence of potential actin manipulation by bacterial flagella. That flagella may be able to plug into this complex system by itself in standard conditions is intriguing, given the lifestyles of their associated pathogens. Conversely, there was no evidence that the *in vitro* binding interactions of H7 flagella with CFL1 affected actin polymerisation synergistically. It is possible that additional factors are required for this; if flagella can gain access to the host cytosol directly, the presence of other actin regulators and different micro-environments may inhibit or enhance these interactions to affect host cell binding or invasion.

In summary, the results of this study demonstrate that *S.* Typhimurium and *E. coli* O157:H7 flagella become intimately associated with host cell membranes during initial adherence. This association either results in or is a consequence of host membrane deformation that may cause disruption, with any disruption likely to be a consequence of flagella rotation. *S.* Typhimurium and *E. coli* O157:H7 can bind to actin and actin-binding proteins. These interactions can influence cytoskeletal dynamics *in vitro* and this may be relevant during bacterial colonisation. An understanding of how connected these two phenomena are will require further work to elucidate the molecular mechanisms involved.

## Materials and Methods

### Bacterial strains and growth conditions

Bacterial strains (Table 1) were stored as saturated cultures in Lysogeny Broth (LB) broth with 25% glycerol at −70°C. Bacteria were grown in LB broth at 28°C-30°C (*E. coli* O157:H7) or 37°C (*S*. Typhimurium) at 200 rpm. Antibiotics when required were chloramphenicol or kanamycin at 50 μg/ml and ampicillin at 100 μg/ml. To assess strain motility, fresh colonies were stab-inoculated once into 0.3% (w/v) LB agar. Inoculated plates were incubated at room temperature (RT) for 36 h. Assays were carried out in quadruplicate from separate colonies.

### Mutant strain construction

The flagellin gene *fliC_H7_* from the *E. coli* O157:H7 strain TUV93-0 was exchanged with *fliC_H6_* according to the allelic exchange method published by Blomfield *et al.* (55). Briefly, homologous recombination of pEBW6 or pEBW7 (Table S1) with TUVΔ*fliC* (Table 1) and counter-selection of *sacB* with sucrose was used to generate TUVΔ*fliC* chromosomally complemented with *fliC_H6_* (*TUVfliC*_*H6F*_) and *fliC_H7_* (TUV*fliC*_*H7F*_) respectively. Strains were validated by PCR with *sacB*, *fliC* locus and O157 specific primer pairs (Table S2), Sanger sequencing, motility in 0.3% (w/v) LB agar and wide-field fluorescence microscopy of saturated cultures with 1:1000 dilutions of H6- and H7-specific rabbit IgG (Table S3). The TUV93-0 *motA* mutant (ZAP1575, Table 1) was constructed by allelic exchange using pTOF25*motA* (Table S1) and primer sets: No-motA, Ni-motA, Co-motA and Ci-motA (Table S2). The deleted region corresponds to 2653444-2654278 of EDL933 and was verified by sequencing. A chromosomally complemented strain was made (ZAP1576, Table 1) using the amplicon of primers No-motA/Co-motA (Table S2) and cloning and exchange with the resultant pTOF25 construct. This *motA* complemented version of ZAP1575 has restored motility (Fig. S3). P22 transduction of phase-locked (*hin*) and *motA* mutations into the SL1344 background from the LT2 strains (Table 1) was carried out using standard transduction protocols from strains used and verified in previous studies (Table 1).

### Antibodies and purified proteins

Abbreviations and/or Uniprot accession codes are in brackets. Sources, details and use of antibodies and stains are described in Table S3. Rabbit skeletal muscle αβ-actin (ACTB, P29751), human platelet βγ-actin (ACTB, P60709), recombinant human cofilin-1 (CFL1 P23528) and arp2/3 complex (ARPC4 Q148J6) from bovine brain were all purchased from Cytoskeleton Inc. Recombinant human galectin-4 (GAL4 P56470) and gelsolin (GSN Q3SX14) from bovine plasma were purchased from R&D Systems and Sigma respectively.

Flagella were sheared from mutants TUV*fliC_H7F_* (H7, Q7DBI0), TUV*fliC_H6F_* (H6, B7USU2), *TUVfliC*∷pEW5 (H48, P04949), *SL1344*Δ*fljB_P2_* (phase-1, P1, FliC E1WGJ5) and SL1344Δ*fliC_P1_* (phase-2, P2, FljB E1WA22), using an adapted protocol (56). TUV93-0 derivatives were cultured at 30°C and SL1344 derivatives at 37°C. Briefly, strains were cultured on 0.3% (w/v) LB agar for 24 h. LB was inoculated with 10 μl agar plugs from the leading edge of the motility halo and incubated at 200 rpm for 16 h. For purification of H48 flagella, *TUVfliC^−^*∷pEW5 was grown in the presence of ampicillin at all times. *TUVfliC^−^*∷pEW5 was further sub-cultured in LB in the same conditions to OD_600_ 0.6, at which point 1 mM IPTG was added and cultures were then grown to OD_600_ 1-1.5. All cultures were centrifuged at 4,100 × *g* at 4°C for 30 min and pellets were re-suspended 1:10 initial culture volume with cold Dulbecco A phosphate-buffered saline (PBS). These were then sheared for 2 min on ice with at maximum speed with an IKA T-10 homogeniser (Ultra-Turrax). Two to five rounds of centrifugation at 4,100 × *g* at 4°C for 15 min, then centrifugation once at 16,000 × *g* at 4°C for 10 min were undertaken to obtain bacteria-free supernatants. These were centrifuged at 145,000 × *g* 4°C for 1.5 h and re-suspended in PBS (or 50 mM Tris-HCl pH7.5 for actin polymerisation assays) at 1:500 of initial culture volumes. Protein concentration and purity was assessed using BCA, then confirmed by coomassie staining (Imperial protein stain, Thermo Fischer) and densitometry after SDS-PAGE.

Sheared flagella were monomerised according to the protocol published by Smith *et al.* (57). Flagella preparations were incubated at 70°C for 15 min then filtered by centrifugation at 5000 × *g* at 4°C for 30 min using 100 kDa 4 ml filter units (Millipore). Filtrates were confirmed by size-exclusion HPLC to be exclusively monomeric.

### Tissue culture

All cells were maintained at 37°C, in 5% CO_2_ and 80% humidity. The embryonic bovine lung epithelial (EBL, laboratory stocks), murine fibroblast (3T3, laboratory stocks) and HEK293 (laboratory stocks) cell lines were maintained in Dulbecco’s minimal essential medium (DMEM, Sigma) supplemented with 10% (v/v) foetal bovine serum (FBS, Sigma), 1 U of penicillin (Invitrogen), 1 μg/ml of streptomycin (Invitrogen) and 2 mM L-glutamine (Invitrogen) unless otherwise stated. The porcine intestinal epithelial cell line IPEC-J2 was maintained as described by Schierack *et al.* (58), in HAMS/F-12 1:1 with 5% FBS, 1 U of penicillin and 1 μg/ml of streptomycin. Prior to bacterial infection, cultured cells were washed in MEM-HEPES and incubated at 37°C, 5% CO2, 80% humidity for 1-2 h. All bacterial infections were incubated in these conditions for the time specified unless otherwise stated.

### Primary cell culture

Bovine rectal and porcine colonic epithelia were isolated from abattoir-derived tissue as described previously (59). The intestinal tissues were collected from material discarded as part of the normal work of the abattoir; while no licence was required for this, the relevant permissions were obtained from the West Lothian abattoir. The mucosal scrapings from bovine terminal rectal or porcine colonic tissue from a local abattoir were digested in DMEM [1% (v/v) fetal calf serum (FCS), 100 U/ml penicillin, 30 μg/ml streptomycin, 25 μg/ml gentamicin] containing 75 U/ml collagenase and 20 μg/ml dispase (Roche) with gentle shaking at 37°C until isolated crypts could be observed microscopically. Crypt enrichment from undigested, contaminating gut microflora and single cells including fibroblasts, was performed using a series of differential centrifugation steps with DMEM containing 2% (w/v) sorbitol. Approximately 500–700 crypts were seeded per well on to collagen (Nutacon for bovine cultures, Porcogen for porcine cultures) coated plates. The bovine cells were grown for 5 days before use, and the porcine cells were grown to a stage of confluence (approximately 3 × 10^5^ cells per well, typically at 10–14 days following initial primary epithelial cell culture).

### Confocal microscopy

Unless otherwise stated, all samples for confocal microscopy outlined below were washed three times in PBS, fixed for >20 min RT in 4% (w/v) paraformaldehyde then permeabilised for <1 min with 0.1% (v/v) Triton X-100 prior to immediate washing as above and staining as below. All stains and antibodies were diluted in PBS. Where stained, cell nuclei were stained with DAPI (Table S3), and slides or coverslips were mounted with ProLong Gold (Invitrogen) unless otherwise stated. Image data was acquired with a Zeiss Plan Apochromat 1.4 NA x63 oil immersion lens and a multi-track (sequential scan) experimental set up on a Zeiss LSM510, using Axiovision software, unless otherwise stated.

For examination of H7 flagella interactions with actin, primary bovine rectal epithelial and EBL cells were used. For primary epithelial interactions, bovine crypts isolated from rectal epithelia were seeded onto collagen-coated 4-well Thermanox chamber slides as described (see primary cell culture section). Infection was for 1 h with 1×10^7^ *E. coli* O157:H7 TUV93-0 grown at 28°C 200 rpm, sub-cultured to OD_600_ 0.3-0.4 and re-suspended in MEM-HEPES. These were then fixed, permeabilised and stained as described above, except that primary antibodies were labelled with FITC-conjugated α-rabbit IgG (Sigma). F-actin was then stained with Texas Red-conjugated phalloidin diluted 1:250, after which cell nuclei were stained with DAPI diluted 1:1000.

Where EBL cells were used, they were seeded onto glass coverslips 24 h prior to infection with exponential-phase *E. coli* O157:H7 TUV93-0. TUV93-0 was grown in 0.3% (w/v) LB agar, sub-cultured into LB broth at 30°C for 16 h 200 rpm, then sub-cultured 1:100 in MEM-HEPES supplemented with 250 nM Fe^2+^ and 0.2% (w/v) glucose at 37°C to an OD_600_ 0.3. EBL cells were infected with TUV93-0 at an MOI of 20 for 3 h. Samples were treated with α-H7 rabbit IgG diluted 1:100, then α-O157 mouse IgG diluted 1:1000, followed by a co-incubation of AlexaFluor_568_-conjugated α-rabbit IgG and AlexaFluor_568_ α-mouse IgG both diluted 1:1000. F-actin was stained with FITC-conjugated phalloidin (Molecular Probes) diluted 1:100 for 2 h. The cell nuclei were stained with DAPI diluted 1:5000.

For investigation of *S*. Typhimurium flagella interactions with host epithelia, confluent IPEC-J2 monolayers on glass coverslips were washed three times with MEM-HEPES immediately prior to infection with *S*. Typhimurium SL1344. Bacterial cultures were incubated in LB with 300 mM NaCl at 37°C 200rpm for 16 h, then diluted to an OD_600_ 0.3 in MEM-HEPES. IPEC-J2 cells were infected at an MOI of 20, for 20 min. Samples were fixed with 2% (w/v) PFA for 20 min then permeabilised with 0.2% (v/v) triton X-100 for 5 min. Samples were treated with P1 or P2 specific rabbit IgG (α-Hi diluted 1:100 and α-H2 diluted 1:500 respectively) then α-O4 rabbit IgG diluted 1:500. This was then labelled with FITC-conjugated α-rabbit IgG diluted 1:1000. F-actin was stained with Texas Red-conjugated phalloidin diluted 1:500 and slides were mounted in VectorShield (Vector Labs). Membrane staining of *S*. Typhimurium infected cells was undertaken in the same way, except that cells were infected for 10 min before fixation, and prior to permeabilisation, membranes were stained with Texas Red-conjugated wheat-germ agglutinin diluted 1:1000 for 2 h (32). Additionally, instead of Texas Red-conjugated phalloidin, F-actin was stained with AlexaFluor_647_-conjugated phalloidin.

For investigation of H7 flagella interactions with host cell membranes, HEK293 cells were transfected with a CRD-STREX-GFP construct that leads to fluorescent staining on the inner side of the plasma membrane (31). Briefly, 5×10^4^ HEK293 cells were seeded onto glass coverslips in DMEM with 10% (v/v) FBS, 100 U/ml penicillin and 30 μg/ml streptomycin, and cultured to 40-70% confluence. 150 ng of CRD-STREX-GFP plasmid was diluted in proprietary EC buffer (Qiagen) and enhancer was added to it at 1:125 (μl/ng DNA) and incubated for 5 min at RT. Effectene (Qiagen) was added in a 3:1 ratio to the enhancer and incubated for 10 min at RT. HEK293 cells were washed with PBS and fresh maintenance medium added. The DNA-Effectene mixture was diluted 1:5 in maintenance medium and added to cells 1:2. Protein expression was determined 24 h post transfection using GFP expression as a read-out. These cells were infected with *E. coli* O157:H7 TUV93-0 for 90 min. Fixed then permeabilised samples were labelled with α-H7 rabbit IgG then α-O157 rabbit IgG (Table S3) diluted 1:1000 and 1:100 respectively. Primary antibodies were labelled with AlexaFluor_568_-conjugated α-rabbit IgG diluted 1:1000. Volumetric 3D reconstructions of de-convolved images were undertaken using Volocity Visualisation software version 6.3.

For all above confocal microscopy, image data was acquired at optimal z-slice sampling rates as determined by Zeiss software, with a 1024 × 1024 pixel image size. Image data was de-convolved using Huygens software (Scientific Volume Imaging, Netherlands). De-convolved models of image data were analysed and re-slices of z-stacks, montages and projection views were created using NIH ImageJ software, with final figures assembled in Adobe Photoshop CS5 and CS6. Co-incidence analysis was undertaken using the 24/10/2004 version of an open-source co-localisation plug-in downloadable from http://rsbweb.nih.gov/ij/plugins/colocalisation.html. Threshold values were set based on z-stack histograms to determine the Gaussian curve of background readings for each channel, and co-incidence was output at a display value of 150. Display values were designated co-incident, not co-localised, as total co-localisation was <50%.

For pre-/post-permeabilisation labelling, collagen (rat tail type 1, Sigma)-coated coverslips were coated in 35% (v/v) ethanol for 4 h at 37°C, washed in PBS then media, then seeded with IPEC-J2 cells for Salmonella infection and EBL cells for *E. coli* O157:H7 infection 24 h prior to infection. Cells at 50-80% confluence were washed with PBS, then incubated with MEM-HEPES for 1-2h prior to infection with 1×10^7^ cells of mid-exponential phase *S.* Typhimurium SL1344∷pAJR145 (constitutively expressing eGFP) for 20 min, or *E. coli* O157:H7 TUV93-0∷pAJR145 (constitutively expressing eGFP) for 1h respectively. Once infections had taken place, cells were fixed in 4% (v/v) PFA in PBS for 20 min RT°C and blocked with 3% (w/v) BSA (PBS). All incubations were 1h, static, in the dark and at RT°C. All washes and reagents were diluted in PBS alone, and washed 3 times with PBS between steps. Flagella were labelled 1:500 with α-P1+P2 and α-H7 polyclonal rabbit IgG (Table S3) for SL1344 and TUV93-0 infections respectively, then labelled 1:1000 with TRITC conjugated α-rabbit IgG, before washing and permeabilisation with 0.1% (v/v) TX100 for 30s. Flagella labelling was then repeated as above, but with FITC-conjugated α-rabbit IgG, then stained with Alexa-Fluor647 conjugated phalloidin (Invitrogen, 1:125 for 20 min). Coverslips were mounted and images acquired using a Leica HCX Plan Apochromat 1.4 NA x100 oil immersion lens in a multi-track (sequential scan) experimental set up on a Leica SP5II confocal laser scanning microscope with Leica Application Suite X software.

### Correlative transmission electron tomography

IPEC-J2 cells were seeded in rat-tail type I collagen(Sigma)-coated glass-bottomed 35mm μ-Dishes etched with 4 reference grids (IBIDI) 24h before infection with *S.* Typhimurium SL1344∷ pAJR145 (constitutively expressing eGFP) in the conditions stated for pre- and post-permeabilisation labelling experiments. After 20 min infection in the above conditions, cells were washed three times in PBS and fixed in 4% (w/v) PFA. Cells were labelled 1:500 with α-FliC rabbit polyclonal IgG, then 1:1000 with FITC-conjugated α-rabbit IgG (Table S3). Dishes were not mounted but instead imaged in PBS using a Leica SP5II confocal laser scanning microscope to determine locations of cell-associated flagellated bacteria on the 4 reference grids.

Cells were fixed and embedded in preparation for electron tomography as follows. All wash steps and treatments were diluted in ddH_2_O, unless otherwise stated. Fixation was undertaken in 2.5% (w/v) glutaraldehyde (100mM sodium cacolydate pH 7.2) for 20 min, then washed three times before post-fixing with 1% (w/v) osmium tetroxide for 20 min. After a further three washes, cells were then stained with 3% (w/v) uranyl acetate for 20 min and washed three more times. Cells were then dehydrated with graded ethanol washes for 5 min each at 70%, 80%, 90%, 96%, 100% (v/v) before adding 1:1 epon epoxy resin for 1 h 30 min, rocking, to embed. Embedded samples were then baked at 200°C for 5 d before removal from glass dishes with snap-freeze/thawing. The reference grids on the remaining resin were then mounted onto stubs, trimmed to areas of interest determined by confocal microscopy (see also Olmos et al., (60)), and cut into 300 nm thick sections using a Leica EM UC7 with an IC80HD camera. Serial sections of areas of interest were placed onto film-coated copper slot grids (Agar Scientific) and further stained with 3% (w/v) uranyl acetate for 20 min to enhance contrast before imaging. Washed grids were labelled with 15 nm gold particles both to the top and bottom of sections for 5 min then blotted dry.

Samples were mapped to determine the location of flagellated and cell-associated bacteria using a FEI 120kV BioTwin Spirit Transmission Electron microscope. Then TEM tilt series’ (between −60° to +60 in 1.5° increments) were acquired using a FEI 200kV Twin Lens Scanning Transmission Electron microscope and Xplore3D software (FEI). The tilt series’ were reconstructed into electron tomograms using the standard workflow of IMOD and 3dMOD software packages (Boulder Laboratory for 3-D EM of cells, University of Colorado, USA), then analysed and presented using Amira 6.0.0, where an anisotropic diffusion filter was applied to the data, and FIJI software.

### Immuno-gold staining and scanning-electron microscopy

The ultra-structural details of flagella interaction with specified epithelial cell types were visualised using Hitachi 4700 Field Emission Scanning Electron microscope. The specimens were fixed in 3% gluteraldehyde in 100 mM sodium cacodylate buffer (pH 7.4) and processed without permeabilisation for SEM as described previously (59). Immuno-gold labelling was visualised by detecting back-scattered electrons; overlays of back scattering and secondary electrons were false-coloured in Adobe Photoshop CS4 for contrast.

### Haemolysis assays

All bacterial cultures were inoculated with agar plugs from the leading edge of the motility halo of fresh motility plates of the strains indicated. These cultures were incubated statically in LB at RT for 16 h. Cultures were adjusted to equal OD_600_ before addition to a V-bottomed 96 well plate 1:1 (50 μl each) with 50% (v/v) washed sheep red blood cells (RBCs, Oxoid) in PBS. The bacteria were driven into contact with the RBCs by centrifugation at 2000 × *g* for 5 min RT, then gently mixed by pipetting. Plates were incubated 37°C 2 h. Samples were diluted 1:1.5 with 150 μl PBS then gently mixed by pipetting. Plates were then centrifuged 2000 × *g* for 10 min RT to pellet RBCs. The A_405_ of 100 μl of supernatants were then measured as a read-out of total haem release. Independent replicates were undertaken 3-5 times, with technical replicates of 3-4. The mean level of haem release caused by incubation with LB alone was subtracted and data were normalised as % of wild-type haem release. After tests for equal variance were performed on raw data for each strain set in Minitab 17, homoscedastic 2-tailed T-tests were performed on the raw data, between mutant and wild-type strains.

### Preparation of cell lysates

Bovine primary rectal epithelial cells were harvested by washing twice in PBS followed by incubation with TripLE Express (Gibco) at 37°C, 5% CO_2_, 80% humidity for 10 min. An equal volume of PBS was added and then cells were scraped off, collected and centrifuged at 300 × *g* for 2 min at RT. Cells were washed twice by re-suspending cell pellets and centrifuging in PBS or Hank’s Balanced Salt Solution (HBSS). The cell pellet was re-suspended in PBS or HBSS, subjected to 5 cycles of snap-freezing in ethanol and dry-ice and thawing at RT, and clarified by centrifugation at 18,000 × *g* for 10 min at 4°C.

### Western and far-Western blotting

80 μg of primary epithelial cell lysates or 1 μg purified receptor candidates separated in SDS-PAGE gels were Western-blotted onto nitrocellulose (Amersham, GE Healthcare) with Schaeffer-Nielsen buffer (48mM Tris, 39mM glycine, 20% (v/v) methanol, 0.04% (v/v) SDS, pH 9.2), at 15V for 30-60 min. Blots were blocked in Carbofree (Vector Labs) or 0.1% (v/v) Tween_20_ (Sigma) in PBS (PBST) with 5% (w/v) skimmed milk powder (Sigma, Marvel), at RT for 2 h or 4°C for 16 h. Blots were washed three times between all subsequent steps in PBST alone for 15 min at RT, with rocking. For both Western and far-Western blots, antibodies were diluted in 1% (w/v) skimmed milk powder in PBST and incubated with blots for 1 h, rocking at RT.

Prior to antibody labelling, far-Western blots were first probed with 1 μg/ml H7 flagella in PBS for 3 h, rocking at RT. H7 flagella were then treated with α-H7 rabbit IgG diluted 1:1000, and this was labelled with horseradish peroxidase (HRP)-conjugated α-rabbit IgG diluted 1:1000. Western blots of cell lysates were probed with α-cofilin-1 mouse IgG diluted 1:500 or α-galectin-4 goat IgG diluted 1:1000, then a 1:1000 dilution of HRP-conjugated α-mouse or goat IgG respectively. Blots were then developed using Pico-West SuperSignal ECL reagents (Thermo Fischer) in a G:box (Syngene), and captured using GeneSnap (Syngene).

### Pull-down assays

Cyanogen bromide (CnBr) activated sepharose 4B lyophilised beads (GE Healthcare) were used according to manufacturer’s instructions. 50 μg of H7 flagella was added 2:1 to CnBr beads for 16 h at 4°C, rocking, in coupling buffer (100 mM NaHCO_3_ pH 8.3 containing 500 mM NaCl). Excess H7 flagella were washed off beads by five cycles of centrifugation at 18,000 × *g* for 30 s then re-suspension of beads in equal volumes of coupling buffer. Following one additional centrifugation as above, beads were re-suspended in blocking buffer (100 mM Tris-HCl, 500 mM NaCl, pH 8.0) and incubated for 2 h static at RT. CnBr beads were then washed ten times by centrifugation as above, with alternate cycles of suspension in coupling buffer or wash buffer (100 mM acetic acid, 100 mM sodium acetate, 500 mM NaCl, pH 4.0). On the final centrifugation, beads were suspended in 180 μg BTRE freeze-thawed cell lysate in HBSS and incubated for 16 h at 4°C, rocking.

CnBr beads were centrifuged at 18,000 × *g* for 30 s then washed by three cycles of centrifugation as above and re-suspension in 0.1% (v/v) PBST. Beads were centrifuged as above and eluted by incubation in 2x SDS-PAGE sample buffer (Sigma) for 5 min at 100°C and centrifuged as above. Eluted proteins were then analysed by 12% SDS-PAGE and Imperial protein staining (Thermo Fischer). Bands visible only in the presence of H7 flagella-coated CnBr beads were considered likely ligand candidates and equivalent cell lysate bands were excised and identified by mass spectrometry.

### Mass spectrometry

Mass-spectrometry, tandem mass-spectrometry and peptide mass fingerprinting were undertaken by Kevin McLean the Moredun Research Institute Proteomics Facility, UK. Protein bands from Imperial protein stained SDS-PAGE gels were excised and delivered to the proteomics facility. Here they were trypsinised, enriched, cleaned-up and matrix assisted laser desorption ionisation time of flight (MALDI-TOF MS/MS) was performed on a Bruker UltraflexII according to their established protocols. Spectra were then input into MASCOT peptide mass fingerprinting software (Matrix Science) as monoisotopic 1+ (m/z) ratios using a ± 50 ppm tolerance, to generate protein identities.

Accurate MW determination of H6, H7, H48, P1 and P2 was undertaken with online HPLC-MS in the facilities of Proteomics and Metabolomics at The Roslin Institute, UK. Samples of sheared flagella were diluted to ~1 pmole/μl in 0.1 % (v/v) formic acid and ~20 pmole was applied to a microbore HPLC column (Dionex Acclaim C18, 4.6 mm i.d., 150mm length, 5 μm beads, 120 Å pore size) pre-equilibrated with 0.1 % (v/v) formic acid by use of a Ultimate HPLC system (Dionex). Bound components were eluted with a gradient of 0.1 % (v/v) formic acid in acetonitrile into the electrospray source of an amaZon ETD ion trap mass spectrometer (Bruker Daltonics, Germany). The mass spectrometer acquired full scan mass spectra with final spectra being an average of 8 trap fills of a maximum averaging time of 200 ms. Signals corresponding to intact flagellin were summed, the raw data was smoothed and background subtracted and then de-convoluted by use of the Bruker proprietary algorithm.

### Enzyme-linked immuno-sorbent assays (ELISA)

1 μg per well of 95% pure human platelet βγ-actin, recombinant human galectin-4 and recombinant human cofilin-1 were adsorbed to 96-well Nunc Maxisorb plates in 100 mM sodium bicarbonate pH 9.6 for 16 h at 4°C. Wells were blocked using Carbo-free (VectorLabs) for 2 h, at RT, before washing. All wash steps involved washing three times in PBST (0.1% Tween_20_). Polymers and monomers of each flagella type (as verified by size exclusion chromatography), were diluted in PBS only at 1000, 500, 100 and 0 ng per well, in specific wells for 3 h, before washing. Antibodies were diluted in PBST and incubated in wells for 1 h, before washing. H6, H7, P1 and P2 flagella specific rabbit IgG (α-H6 and α-H7 1:1000; α-Hi 1:100; α-H2 1:500 respectively, Table S3) followed by HRP-conjugated α-rabbit IgG were used to label flagella-specific binding.

Flagella binding was detected using Pico-West SuperSignal ECL reagents (Thermo Fischer) in a G:Box using GeneSnap (Syngene) at a fixed distance of 575 mm. Densitometry on wells was performed in GeneTools (Syngene), using a fixed spot radius of 25. Data were normalised according to the formula: 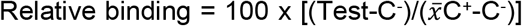, where C^−^ = 0 ng flagella protein, C^+^ = positive detection control. This normalisation takes into account background from antibodies and assay to assay variation. In addition to this, no primary antibody and no ligand coating qualitative controls were performed.

Statistical analysis was carried out in R on log_10_ data from four independent experiments. Data residuals were assessed for normality using a normality plot. As it was not biologically appropriate to statistically compare different flagella types or ligand types, separate linear mixed effects models of relative binding were used to assess the statistical significance of each flagella type binding to each ligand candidate (galectin-4, cofilin-1 and actin). Flagella concentration (1000, 500, 100 ng/well) and polymerisation status (polymer and monomer) were fixed effects and experiment was a random effect. Pairwise comparisons of the interaction between polymerisation statuses for each flagella type in each model were then performed on the data. To correct for the large number of comparisons, an alpha level of p<0.01 was taken as significant.

### Actin polymerisation assays

Actin polymerisation assays were carried out as described by Sitthidet *et al.* with minor modifications (61) and informed by Van Troys *et al.* (37). G-actin was prepared by re-suspending lyophilised pyrene-conjugated rabbit skeletal muscle αβ-actin in G-buffer (10 mM Tris-HCl, 200 μM CaCl_2_, 200 μM ATP, 1 mM DTT, pH 7.5) for 1 h on ice, in the dark. This was then centrifuged at 100,000 × *g* for 2 h at 4°C. The top 80% supernatant was kept on ice in the dark. Test proteins were re-suspended as purified or dialysed with U-tube concentrators (Novagen) into 50 mM Tris-HCl pH 7.5. Molar concentrations of all protein components were calculated using a Bradford assay with a BSA standard curve. Densitometry of coomassie-stained SDS-PAGE gels was used to adjust flagella values.

Assays were carried out in black opaque 96-well plates (Nunc). 1 μM actin was added to wells with 50 mM Tris-HCl pH 7.5. Cofilin-1 and serially diluted flagella preparations were then added, followed by polymerisation buffer (at a final concentration of 5 mM KCl, 0.2 mM MgCl_2_ and 0.1 mM ATP). Samples were excited at 365 nm, and emission data was measured at 407 nm for 1 h, at 30 s intervals. A daily gain value was applied to all wells to normalise actin polymerisation capability and this was calculated using pyrene-actin fluorescence after 1 h polymerisation at RT as above.

Maximum velocity (V_max_) of actin polymerisation was calculated with the formula V_max_ = (A_2_−A_1_)/(T_2_−T_1_), where A = absorbance at 407 nm and T = time. Differences due to addition of flagella were assessed for statistical significance using GLMs in Minitab 16.2.4. In these models, V_max_ was the response and replicate was a random factor. With flagella titration experiments, GLM analysis was carried out on data from 5 independent experiments; flagella type and concentration were fixed factors. Post-hoc Tukey pairwise comparisons of the interaction between flagella type and concentration were carried out. With flagella ± cofilin-1 experiments, GLM analysis was carried out on data from three independent experiments; flagella type and cofilin-1 concentration were fixed factors. Post-hoc Tukey pairwise comparisons of the interaction between cofilin-1 concentration and flagella type were carried out as a very conservative estimate of differences based on the set of assumptions in a GLM statistical model. Therefore, if p-values are <0.05 they are very likely to be valid and so were taken as statistically significant.

### Size exclusion chromatography

Samples were applied to an HPLC column (Sigma TSK G4000SWXL, 4.6 mm i.d., 300 mm length, 8 μm beads, 450 Å pore size) pre-equilibrated with 100 mM Tris-HCl, 200 mM NaCl, 1 mM DTT, pH 8.0, by use of an Ultimate HPLC system (Dionex), measuring absorbance at 220 nm. Columns were calibrated by the maximum peak of elution of individual molecular weight standards: Blue Dextran (2.5 MDa) at ~5 ml, BSA (66 kDa) at ~10 ml and Ribonuclease A (13.5 kDa) and ATP (0.5 Da) both at ~12 ml.

For verification of flagellin polymerisation status, polymeric and monomeric H6, H7, H48, P1 and P2 flagellin purified into 50 mM Tris-HCl pH 7.5 were applied neat to the column three times at 30 min intervals, with three blank runs in between flagella types. Data was analysed by establishing the lowest value of all runs as a baseline and calculating the area under curve (AUC) for each run, and looking at the ratio of filaments (4.5-9 ml ≈ >2MDa- ~300 kDa) to monomers (9-11.25 ml < ~300 kDa) with the formula AUC = [(A_1_+A_2_)/2] × (T_2_−T_1_), where A = absorbance at 220 nm and T = time (ml).

## Acknowledgements

We would like to thank Sutherland Maciver (Edinburgh) for advice on cofilin-1, Kevin Mclean at the Moredun Research Institute proteomics facility for peptide mass fingerprinting, Mike Shipston (Edinburgh) for supplying the STREX-GFP construct, and Phillip Aldridge (Newcastle), Roberto La Ragione (Surrey) and Mark Jepson (Bristol) for supplying reagents and strains, and the Centre for Integrative Physiology (University of Edinburgh) and Wolfson Bioimaging Facility (University of Bristol). We would also like to thank Ian Thompson for his advice and support over the last decade.

## Supplementary Information

**Figure S1.**
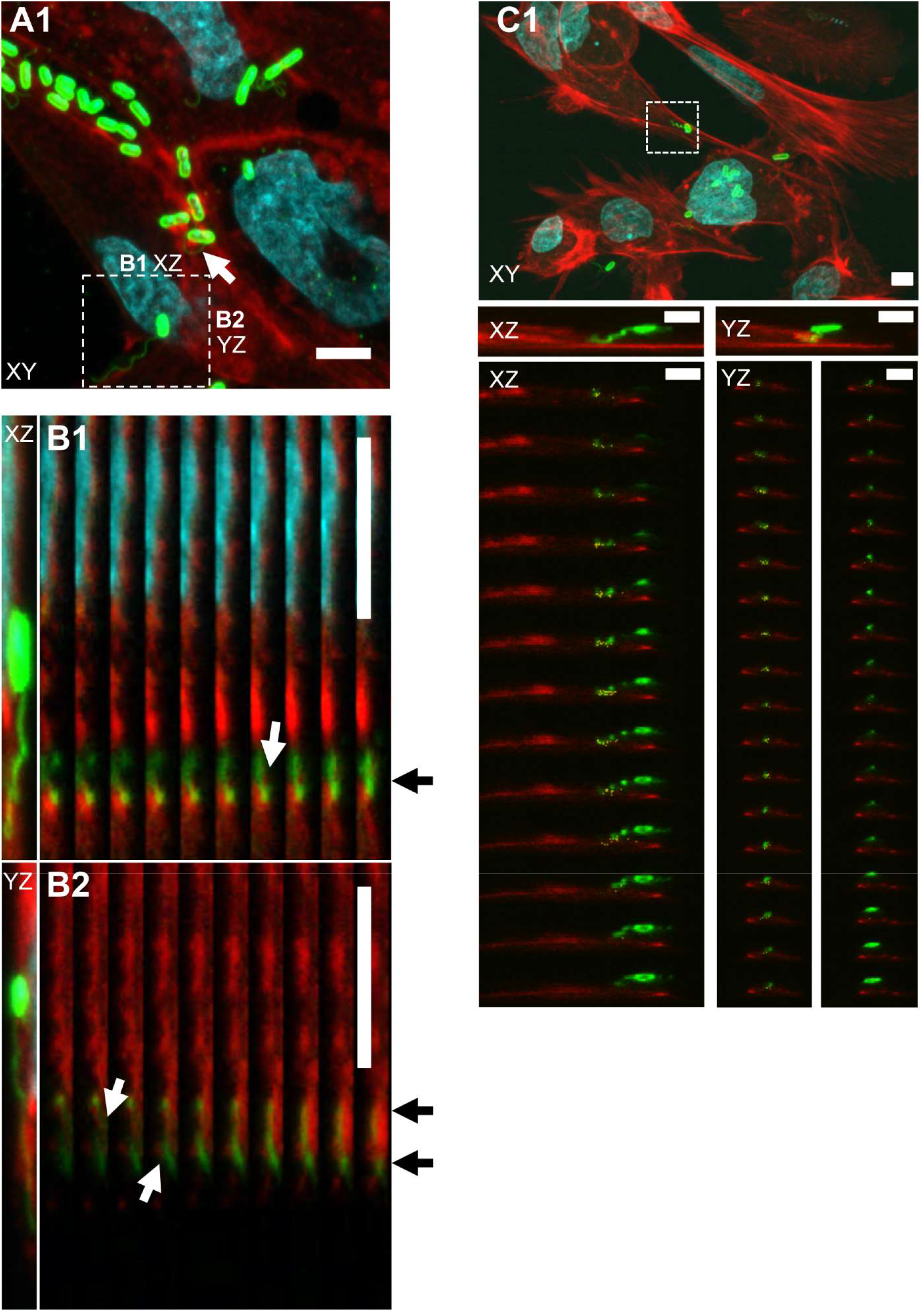
Confocal microscopy of intracellular interactions of H7 flagella with host cells. Confocal z-slices of E. coli O157:H7 TUV93-0 flagella (green) interacting with actin (red). Nuclei staining is cyan. **(A1)** XY projection of TUV93-0 interacting with EBL cells, 3 h post infection. The micro-colony indicated with an arrow shows an H7 flagellum curling round A/E lesions. The labelled inset marks the XZ and YZ projections analysed in B1-B2. **(B1-B2)** Arrows point to specific regions within individual XZ **(B1)** and YZ **(B2)** slices in which H7 flagellum is inside a region of actin staining. **(C1)** Top panel shows an XY projection of the whole field from which the images in Fig. 1A were taken. Middle panels show Fig. 1A XZ and YZ projections, free of actin/flagellum co-incidence labelling. Lower panels show actin-H7 flagellum co-incidence (yellow) across individual z-slices in XZ and YZ planes. Scale bars = 5 μm.

**Figure S2.**
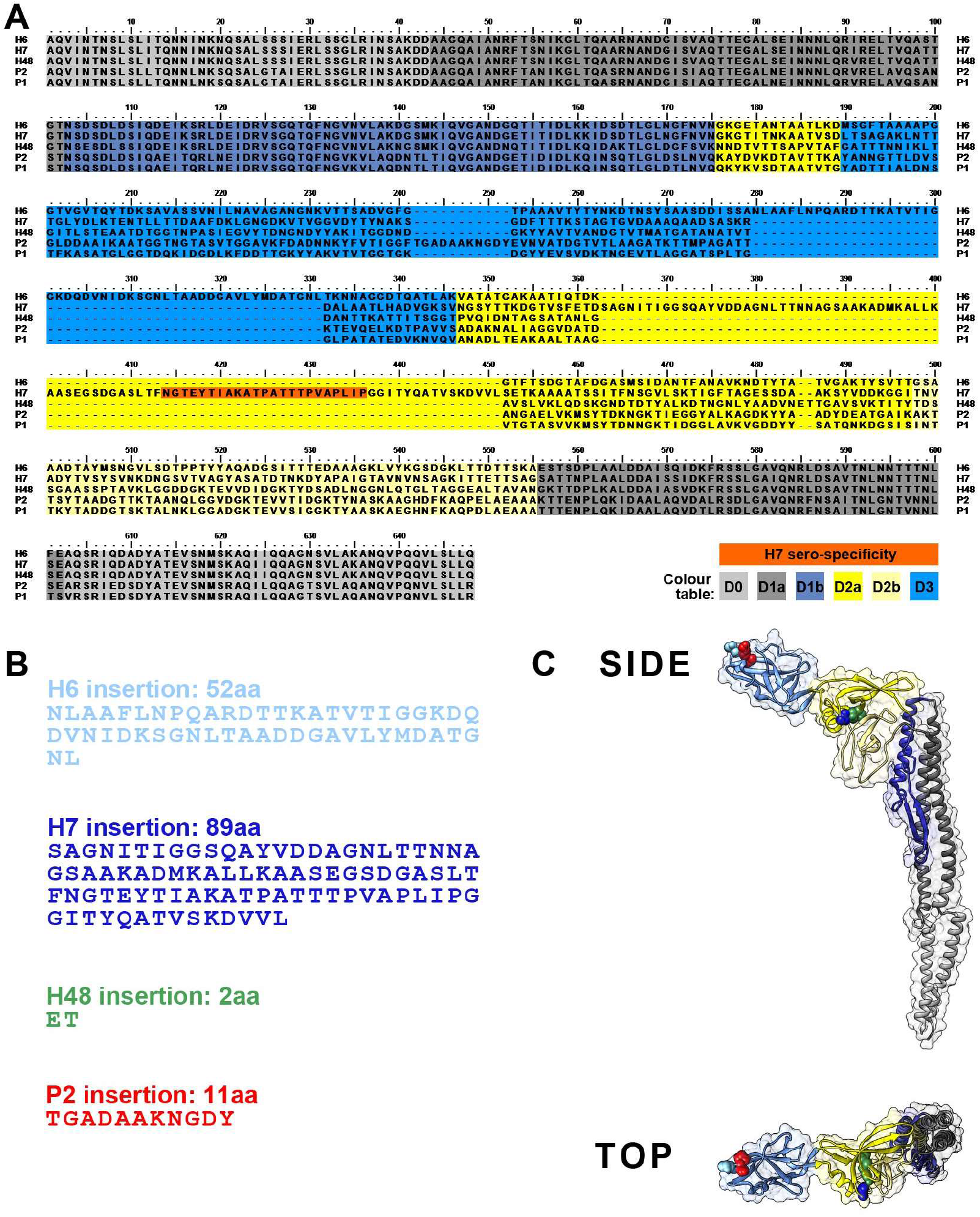
Protein sequence and structural differences between H6, H7, H48, P1 and P2 flagellins. **(A)** Pairwise structural alignments with P1 flagellin (PDB: 1UCU) were generated using the FFAS03 server from its PDB database. Alignments were then stitched together and presented in BioEdit v 7.2.0. Amino acids are coloured by structural domain, as defined by Yonekura *et al*. (9) and outlined in the colour table. This model is partially validated by the presence of the H7-serospecific region in the H7-specific structural insertion ((69), orange). **(B)** Amino acid sequences modelled to be structurally dissimilar to the P1 structural template, as determined by the alignment in (A). **(C)** Location of structural insertions in P1 flagellin indicated by spherically presented side-chains, colour-coded as in (B). The P1 structural model is presented coloured by structural domain as in (A), with top and side views, in USCF Chimera v 1.8 (70).

**Figure S3.**
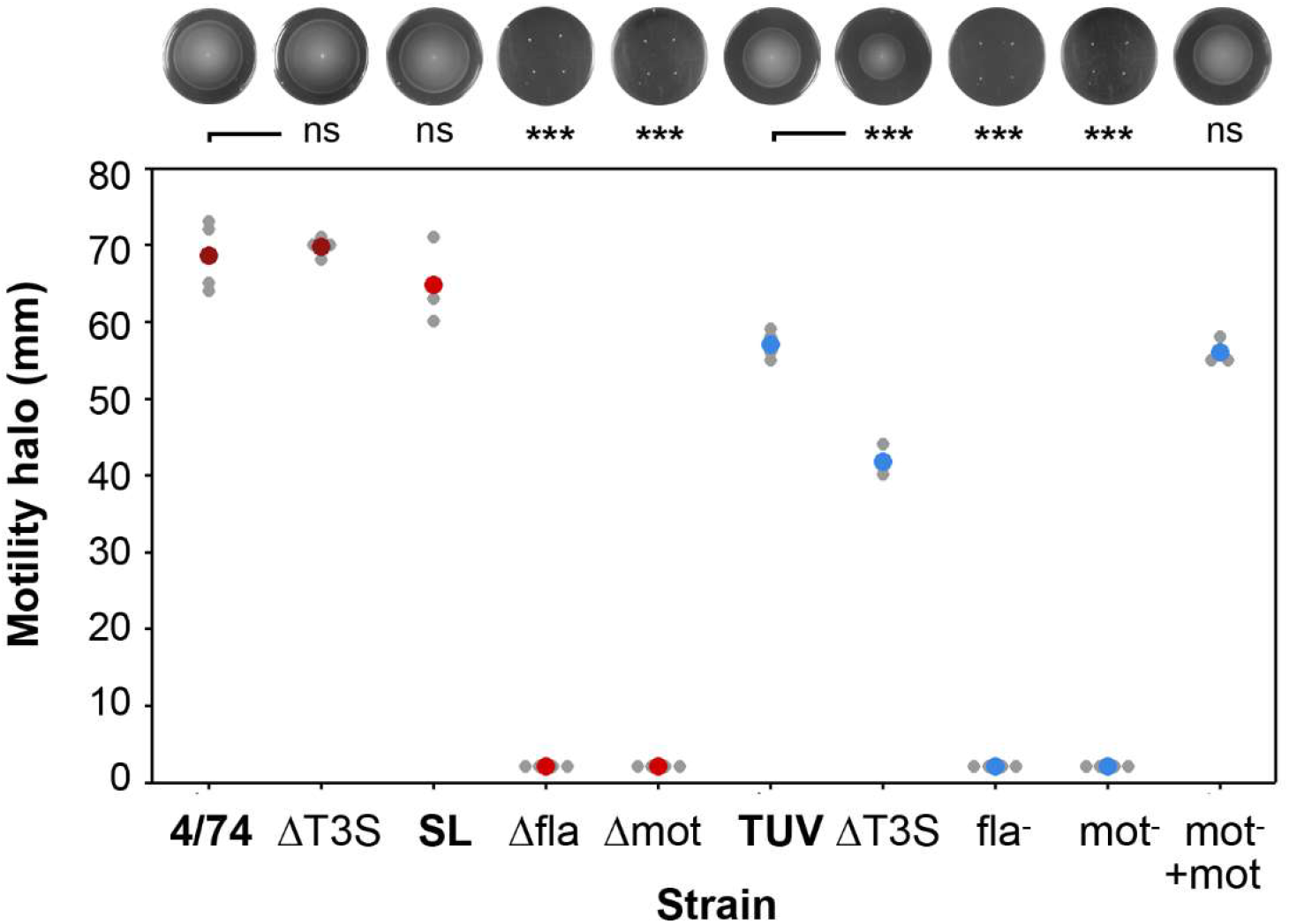
Motility *S.* Typhimurium and *E. coli* O157:H7 strains used in haemolysis assays. Motility of strains used in Fig. 6 was measured by the radius of growth after inoculation into 0.3% (w/v) LB agar at RT for 36 h. SL= SL1344. The top panel shows representative motility plates, and the bottom panel shows radii measurements from point of inoculation of four biological replicates. Statistical analyses of these are presented as 2-tailed homoscedastic students T-tests (p≤0.0001 = ***) against WT (4/74 or TUV93-0).

**Figure S4.**
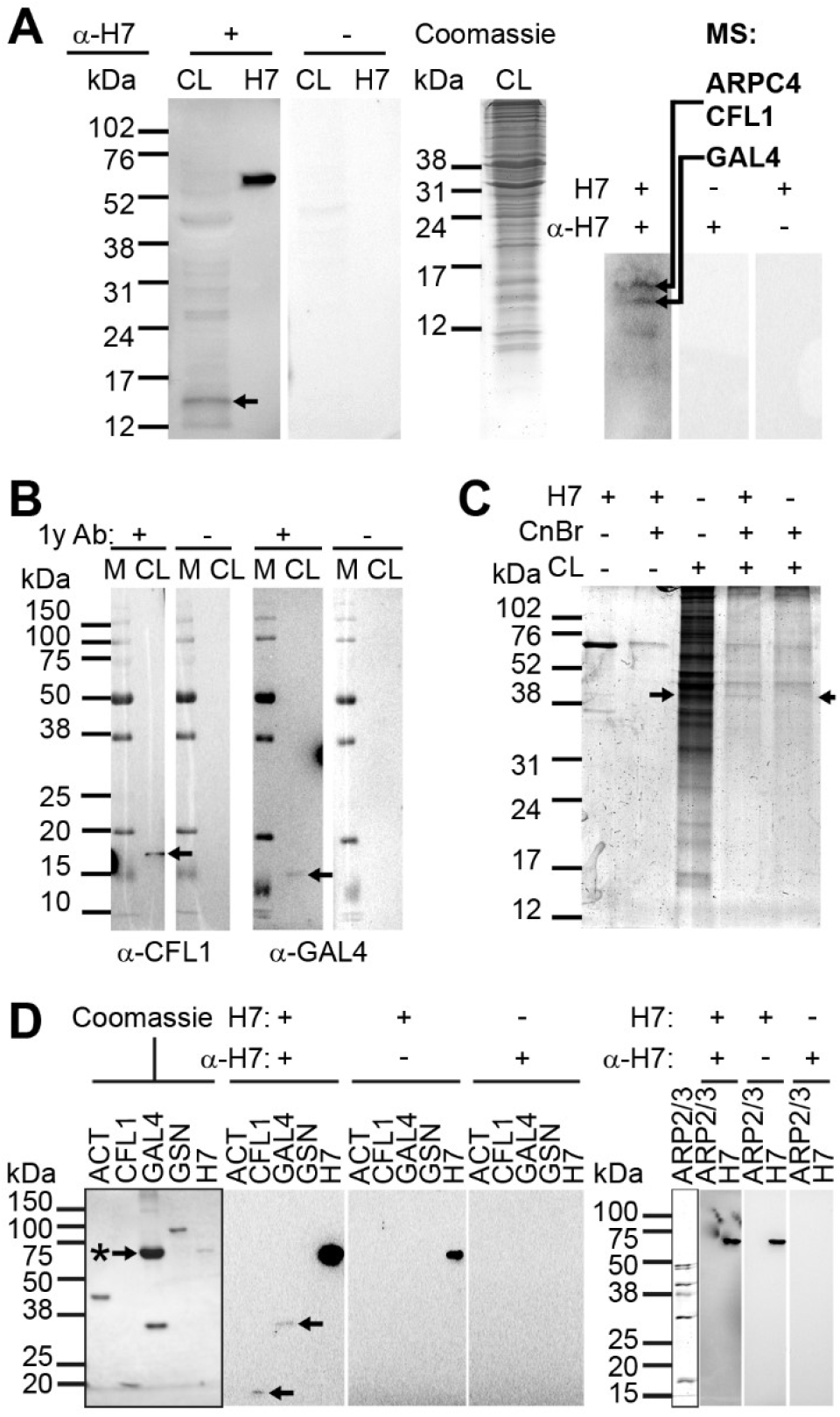
Binding substrates for H7 flagella present in bovine terminal rectum epithelial lysates. **(A)** Far-Western blot of primary bovine rectal epithelial cell lysates (CL) probed with H7 flagella and H7-specific antibodies. Left-hand blots show one reacting band (arrow) resolved further (right) and the bands indicated were identified as Arp2/3 complex sub-unit 4 (ARPC4), cofilin-1 (CFL1) and galectin-4 (GAL4) by mass-spectrometry (MS). **(B)** Western-blots of primary bovine rectal epithelial cell lysates (CL) confirmed the presence of CFL1 and GAL4 between 15-20 kDa with α-CFL1 and α-GAL4 antibodies (arrows, Table S3). The size discrepancy of GAL4 can be explained by its degradation or processing within cell lysates, as has been observed previously (71, 72). **(C)** Pull-down of bovine β-actin from primary bovine rectal epithelial cell lysates (CL) by H7 flagella cross-linked to CnBr-activated Sepharose beads. The cell lysates were prepared by freeze-thawing. Pre-cleared empty beads were used as a negative control (final lane). Arrows indicate 38-50 kDa protein bands in the coomassie-stained gel that were excised and identified as β-Actin (ACTB1) by MS. **(D)** Far-Western blots of 1 μg purified human βγ-actin (ACT), recombinant human CFL1, recombinant human GAL4 and purified gelsolin (GSN) from bovine plasma with negative control blots on the left, and 1 μg purified ARPC4 from bovine brain with detection controls on the right, probed with H7 flagella and H7-specific antibodies as indicated. The ~60 kDa band indicated with an arrow marked with an (*) is 0.1% (w/v) BSA, a carrier protein for GAL4. 0.5 μg H7 was loaded as a positive detection control. Detected bands are indicated with arrows. Marker lane (M) sizes are in kDa on the left throughout.

**Movie S1. Co-localisation of *E. coli* O157:H7 flagella with phalloidin on a bovine terminal rectal epithelial cell.** A rotating three-dimensional projection of a confocal micrograph that has captured *E. coli* O157:H7 (green) flagella coincident with bovine primary rectal epithelial actin (red, Fig. 1A). Co-incident staining of actin and O157:H7 is shown in yellow. The 3D projection was made and presented using NIH ImageJ software.

**Movie S2. Tomographic slice of the flagellated *S.* Typhimurium within an epithelial cell.** A view up and down through the 3D projection of the tomogramic slice shown in figure 5A1, zooming in to the inset shown in figure 5A2. An anisotropic diffusion filter was applied to reduce noise.

**Movie S3. Tomographic slice of the flagellated *S.* Typhimurium inside a membrane ruffle.** A view up and down through the 3D projection of the tomogramic slice shown in figure 5B1, zooming in to the inset shown in figure 5B2. An anisotropic diffusion filter was applied to reduce noise.

**Movie S4. Tomographic slice of the flagellated *S.* Typhimurium at the point of induced uptake into an epithelial cell shown in figure 5C**. A view up and down through the 3D projection of the tomogramic slice shown in figure 5C1, with segmentation analysis of host cell membranes shown in red, zooming in to the inset shown in figure 5C2. An anisotropic diffusion filter was applied to reduce noise.

---

**Table S1.**
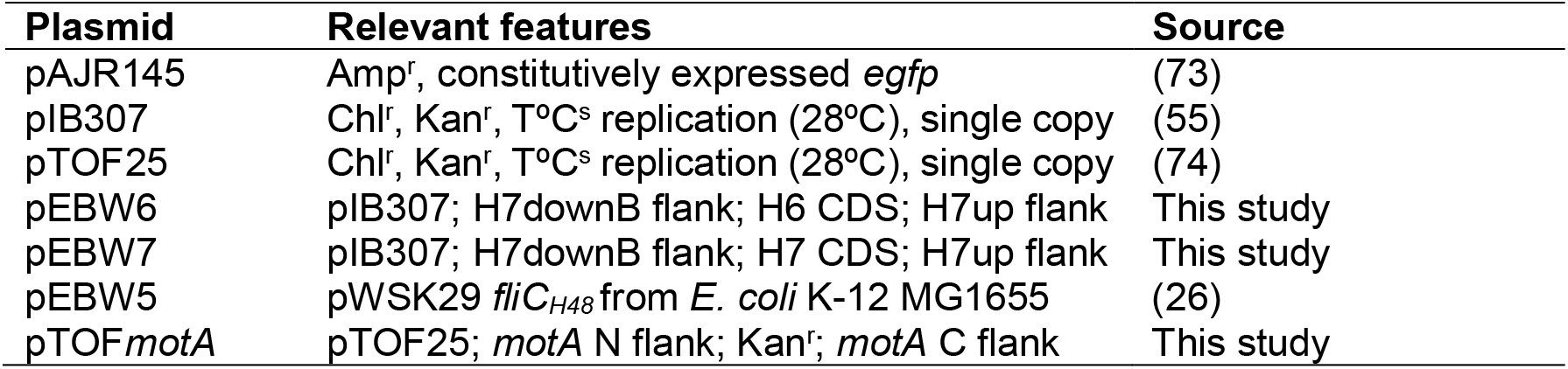
Plasmids used or constructed in this study.

**Table S2.**
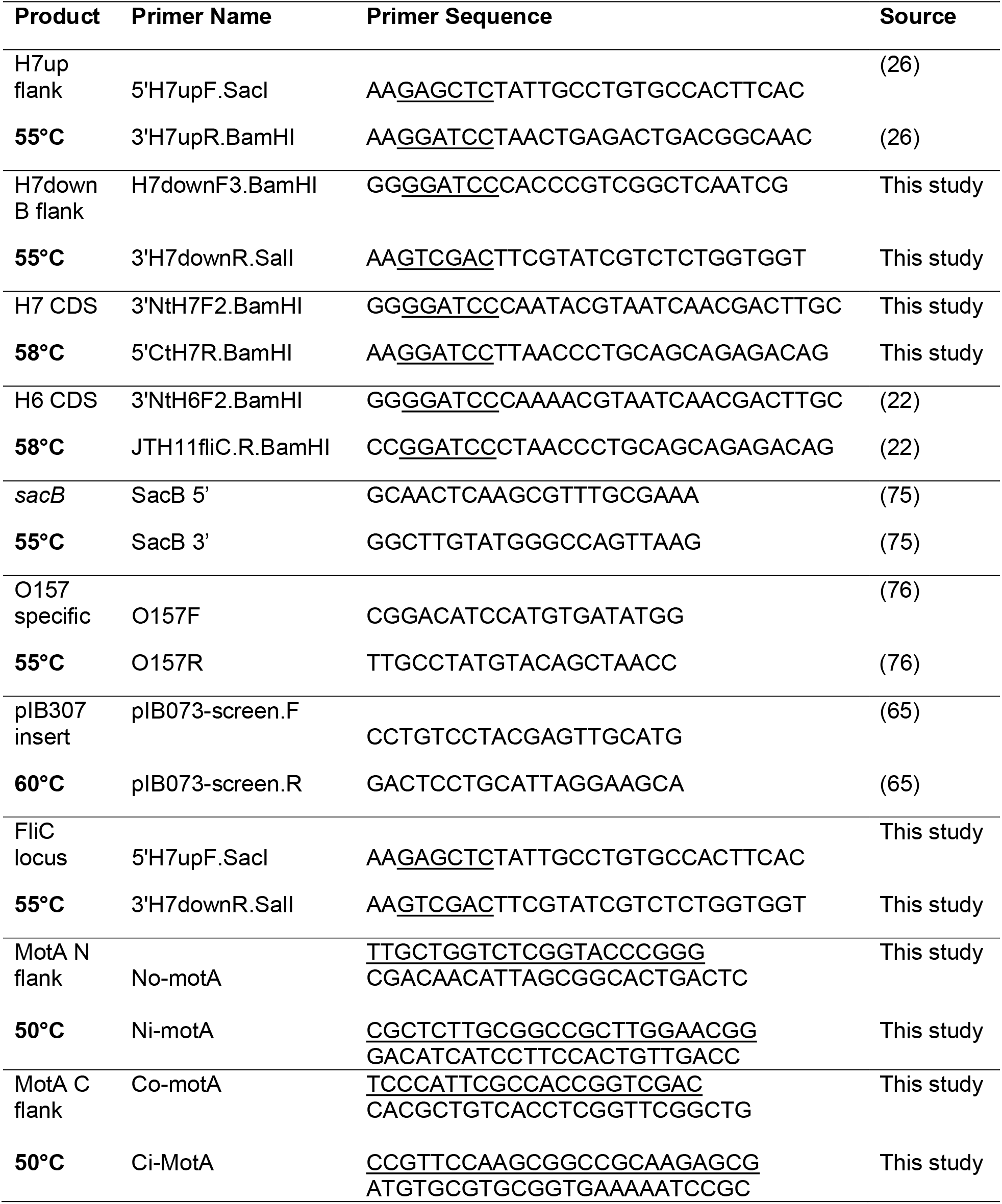
Primer pairs used in this study. Annealing temperatures used indicated in bold.

**Table S3.** Antibodies and stains used in this study.

| **Primary Antibody** | **Details** | **Source** |
| --- | --- | --- |
| α-H6 | polyclonal rabbit IgG | Mast Assure |
| α-H7 | polyclonal rabbit IgG | Mast Assure |
| α-FliC (P1) | polyclonal rabbit IgG | Ariel Blocker Lab,<br>University of Bristol |
| α-Hi (P1) | polyclonal rabbit IgG | Mast Assure |
| α-H2 (P2) | polyclonal rabbit IgG | Difco |
| α-P1+P2 | polyclonal rabbit IgG | Difco |
| α-O157:H7 | polyclonal rabbit IgG | Mast Assure |
| α-O157 | monoclonal mouse IgG | Abcam |
| α-O4 | polyclonal rabbit IgG | Mast Assure |
| α-cofilin-1 | monoclonal mouse IgG | AbDSeroTec |
| α-galectin-4 | polyclonal goat IgG | R&D |
| **Secondary Antibody** | **Details** | **Source** |
| α-rabbit IgG-HRP | polyclonal goat IgG | R&D |
| α-goat IgG-HRP | polyclonal rabbit IgG | R&D |
| α-mouse IgG-HRP | polyclonal donkey IgG | R&D |
| α-rabbit IgG-FITC | polyclonal goat IgG | Sigma |
| α-rabbit IgG-FITC | polyclonal goat IgG | R&D |
| α-rabbit IgG-10nm gold | polyclonal goat IgG | British Biocell Intl. |
| α-rabbit IgG- AlexaFluor568 | polyclonal goat IgG | Molecular probes |
| α-mouse IgG- AlexaFluor568 | polyclonal goat IgG | Molecular probes |
| α-rabbit IgG- AlexaFluor594 | polyclonal goat IgG | Molecular probes |
| **Stains** |  | **Source** |
| Phalloidin-Texas Red |  | Invitrogen |
| Phalloidin-AlexaFluor588 |  | Molecular Probes |
| Phalloidin-AlexaFluor647 |  | Invitrogen |
| Phalloidin-FITC |  | Molecular Probes |
| Phalloidin-TRITC |  | Sigma |
| DAPI |  | Merck |
| Wheat-germ agglutinin-Texas Red |  | Invitrogen |

